# Inference of infectious disease transmission using multiple genomes per host

**DOI:** 10.1101/2023.07.28.550949

**Authors:** Jake Carson, Matt Keeling, David Wyllie, Paolo Ribeca, Xavier Didelot

**Affiliations:** Mathematics Institute, University of Warwick, Coventry CV4 7AL, United Kingdom; School of Life Sciences, University of Warwick, Coventry CV4 7AL, United Kingdom; Zeeman Institute for Systems Biology and Infectious Disease Epidemiology Research (SBIDER), University of Warwick, Coventry CV4 7AL, United Kingdom; UK Health Security Agency, London NW9 5EQ, United Kingdom; Department of Statistics, University of Warwick, Coventry CV4 7AL, United Kingdom

**Keywords:** genomic epidemiology, transmission analysis, infectious disease outbreak, within-host diversity and evolution

## Abstract

In recent times, pathogen genome sequencing has become increasingly used to investigate infectious disease outbreaks. When genomic data is sampled densely enough amongst infected individuals, it can help resolve who infected whom. However, transmission analysis cannot rely solely on a phylogeny of the genomes but must account for the within-host evolution of the pathogen, which blurs the relationship between phylogenetic and transmission trees. When only a single genome is sampled for each host, the uncertainty about who infected whom can be quite high. Consequently, transmission analysis based on multiple genomes of the same pathogen per host has a clear potential for delivering more precise results, even though it is more laborious to achieve. Here we present a new methodology that can use any number of genomes sampled from a set of individuals to reconstruct their transmission network. We use simulated data to show that our method becomes more accurate as more genomes per host are provided, and that it can infer key infectious disease parameters such as the size of the transmission bottleneck, within-host growth rate, basic reproduction number and sampling fraction. We demonstrate the usefulness of our method in applications to real datasets from an outbreak of *Pseudomonas aeruginosa* amongst cystic fibrosis patients and a nosocomial outbreak of *Klebsiella pneumoniae*.

## INTRODUCTION

Pathogen genomic data has transformed our understanding of the epidemiology of infectious diseases, whether they are caused by viruses (Grenfell et al., 2004; Pybus and Rambaut, 2009) or bacteria (Didelot et al., 2012; Gardy and Loman, 2018). Most applications concern large-scale pathogen populations, for example to estimate their demographic history (Pybus et al., 2001; Ho and Shapiro, 2011) or the way that their ancestry relates to features of geography (Lemey et al., 2009; De Maio et al., 2015), epidemiology (Volz et al., 2013; Rasmussen et al., 2014) or host population (Mather et al., 2013; Dearlove et al., 2016). Genomic data can however also be useful to perform much finer inference, down to the level of transmission analysis which attempts to reconstruct who infected whom within an outbreak (Cottam et al., 2008; Jombart et al., 2011). Phylogenetic methods have a long successful history and can reconstruct the genealogy of a set of genomes given their sequences (Yang and Rannala, 2012; Kapli et al., 2020). However, a phylogenetic tree is not identical to a transmission tree (Pybus and Rambaut, 2009; Jombart et al., 2011; Romero-Severson et al., 2014). In particular, the nodes in a phylogenetic tree do not correspond to transmission events, but rather to lineages diverging during the evolutionary process that takes places within a host (Didelot et al., 2016). Several methods have therefore been developed over the past few years specifically aimed at the reconstruction of a transmission tree, including SeqTrack (Jombart et al., 2011), outbreaker (Jombart et al., 2014), beastlier (Hall et al., 2015), SCOTTI (De Maio et al., 2016), phybreak (Klinkenberg et al., 2017) and outbreaker2 (Campbell et al., 2018).

Here we focus on one such method for transmission analysis called TransPhylo, which is based on colouring the branches of a dated phylogeny to reveal the transmission tree (Didelot et al., 2014). There are many software tools that can be used to construct such a dated phylogeny, for example BEAST (Suchard et al., 2018), BEAST2 (Bouckaert et al., 2019), BactDating (Didelot et al., 2018), treedater (Volz and Frost, 2017) and TreeTime (Sagulenko et al., 2018). An advantage of the TransPhylo colouring approach is that it separates the initial phylogenetic reconstruction from its epidemiological interpretation, which improves computational efficiency and therefore scalability (Didelot and Parkhill, 2022). Furthermore, the original TransPhylo model (Didelot et al., 2014) has been extended to deal with both partially sampled and ongoing outbreaks (Didelot et al., 2017). Consequently, TransPhylo is a flexible and versatile software to perform transmission analysis using pathogen genomic data (Didelot et al., 2021).

Following infection, many pathogens evolve within hosts on a time scale that is relevant to transmission analysis (Lieberman et al., 2011; Bryant et al., 2013; Biek et al., 2015; Grote and Earl, 2022). Consequently, when information is available about the within-host pathogen diversity, this can help clarify who infected whom (Didelot et al., 2016; Leitner, 2019). This information can come in two forms: either heterogeneities in the genomic sequencing of a single clinical sample, or genomic sequencing of multiple separate clinical samples.

Genetic heterogeneities within a sample are relatively easy to survey, and a few methods have been developed recently with the specific aim of exploiting this type of data to help infer transmission (De Maio et al., 2018; Wymant et al., 2018; Ortiz et al., 2022). However this approach requires the analysis of short sequencing reads individually which can be difficult and error-prone; additionally the clinical sample may not represent the within-host diversity of the pathogen when it was collected, and it does not contain any information about evolution or changes of diversity over time in the within-host pathogen population.

The alternative approach of sequencing several longitudinal clinical samples is more involved, but can provide a more thorough and reliable overview of the within-host evolution, as exemplified in studies of infection with *Staphylococcus aureus* (Young et al., 2012), *Helicobacter pylori* (Didelot et al., 2013) or *Streptococcus pneumoniae* (Tonkin-Hill et al., 2022). In principle, integrating multiple genomes into a joint model of phylogenetic and transmission trees, such as TransPhylo, is possible by having as many leaves in the phylogenetic tree as there are samples (Didelot et al., 2016; Leitner, 2019). However, this poses a significant number of theoretical challenges to overcome, which is why TransPhylo was not previously able to use more than one genome per host (Didelot et al., 2017; Xu et al., 2020). Here we present a solution to these issues, which leads us to formulate an extended version of the TransPhylo model, inference methodology and software, so that any number of genomes per host can be used as input of a transmission analysis.

## NEW APPROACHES

We extend the latest TransPhylo framework (Didelot et al., 2017) to incorporate multiple samples per host. The model in TransPhylo has three basic ingredients which we detail below, before explaining the changes needed to deal with multiple samples per host. Firstly, a coalescent model with constant population size and temporally offset leaves (Drummond et al., 2002) to represent the within-host evolution. Secondly, a branching process transmission model in which individuals are sampled either once or not at all, so that unsampled individuals can be accounted for in the transmission chains between sampled individuals. Thirdly, a complete transmission bottleneck meaning that only a single lineage is ever transmitted between hosts. In other words the within-host coalescent process is bounded so that the most recent common ancestor within a host occurs after the date of infection (Carson et al., 2022).

The full bottleneck assumption can be problematic in settings where hosts are repeatedly sampled, as the resulting phylogenetic trees may have no compatible transmission trees (Romero-Severson et al., 2014, 2016). Therefore we remove this complete bottleneck assumption, so that the phylogenetic trees are much more likely to have compatible transmission trees. Our main motivation for removing this assumption was to allow for multiple samples per host, but it is also important to note that a number of studies have found that the transmission bottleneck is only partial for many pathogens including HIV (Boeras et al., 2011), FMDV (Cortey et al., 2019), influenza (Ghafari et al., 2020) and *Staphylococcus aureus* (Hall et al., 2019). Relaxing the transmission bottleneck assumption therefore leads to a more generally applicable model, in which it is possible to additionally estimate the scale of the transmission bottleneck.

We also relax the assumption of a constant within-host population size by allowing linear growth, following previous work on HIV (Romero-Severson et al., 2014, 2016; Leitner, 2019). This linear growth model is a generalisation of the constant population size model which can be obtained if the linear growth rate parameter is set to zero. It is also a generalisation of a linear growth with complete transmission bottleneck model (Klinkenberg et al., 2017) since this can be obtained if the linear intersect is zero at the date of infection. The linear growth model therefore has several advantages, on top of being simple and statistically tractable, but other options such as an exponential or logistic growth model could also be used as will be discussed later.

Finally, in the transmission model we add the possibility that hosts are sampled multiple times, while also retaining the possibility that some hosts are sampled only once or not at all. We make the specific choice that the transmission model up to the first sample for each host is exactly the same as previously formulated (Didelot et al., 2017). The times of any further sampling depend only on the first observation times, and not the infection times. Since the infection times and secondary observation times are conditionally independent given the primary observation times, we can infer the infection times without the need to formally define this aspect of the model. In the Methods section we present a full mathematical description of this new extended model, and show how Bayesian inference can be performed using a Markov Chain Monte-Carlo (MCMC) scheme with reversible-jumps (Green, 1995) to accommodate the non-constant dimension of the parameter space.

## RESULTS

### Exemplary analysis of a single simulation

We simulate an outbreak with 100 observed hosts, each with five observations. The observation cut-off time *T* is determined by the simulation in order to return the correct number of observed hosts. The generation time and primary observation time are both Gamma distributed (see section “Epidemiological model” in the Materials and Methods) with shape and scale parameters equal to 2 and 1, respectively. Secondary observations are placed at intervals of 0.25 years following the primary observation. For the transmission model, the offspring distribution is negative binomial with mean equal to the basic reproduction number *R* = 2, and the sampling proportion is *π* = 0.8. The within-host pathogen population size is *κ* + *λτ* at time *τ* after infection, with *κ* = 0.1 and *λ* = 0.2. The resulting simulation contains 124 hosts, four of which are infected with two lineages at the time of infection, one with three lineages, and the remaining 119 with a single lineage.

We investigate the ability of our methodology to recover the model parameters used in the simulation, and to recover transmission links between individuals. We also investigate what benefits are obtained by including multiple observations per host. To this end we construct additional phylogenetic trees by pruning the last observation for each host. Through repetition we obtain phylogenetic trees with four, three, two and one observations per host under the same transmission network. By comparing inference outcomes from these five trees we can establish the extent to which estimates are improved through the inclusion of secondary observations.

We perform 12,000 MCMC iterations for each phylogenetic tree, using the first 2,000 as a burn-in. The prior distribution for *π* is uniform between 0 and 1, and the prior distributions for *R, κ* and *λ* are exponential with mean 1. The posterior means and 95% credible intervals are shown in the Table 1. These results demonstrate that we are able to recover the model parameters used in the simulation, even with no secondary observations. Comparing posterior estimates across the different trees indicates that our estimates of the transmission model parameters *R* and *π* are not considerably improved by the number of secondary observations. This makes sense, as most of the relevant information for these parameters is contained in the primary observation. However, the credible intervals for the coalescent model parameters *κ* and *λ* narrow as more secondary observations are added. Secondary observations provide considerable information about the within-host genomic diversity of infected hosts, leading to more precise estimates.

**Table 1:**
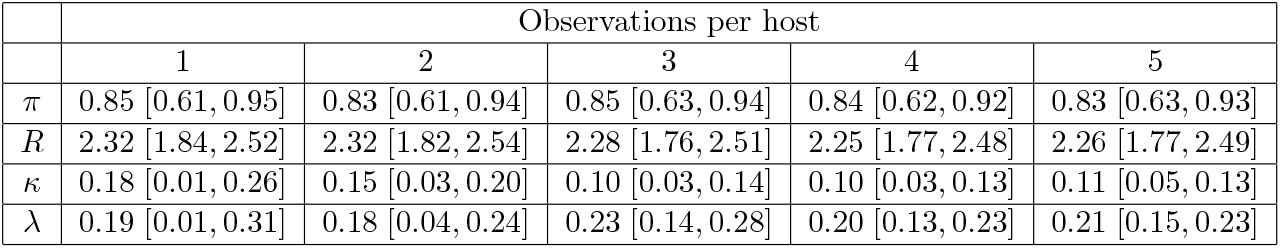
Posterior estimates of the simulation study given as the posterior mean and 95% credible interval. The model parameter is given in the left column, and the remaining columns indicate the number of observations per observed host. The values used in the simulation are *π* = 0.8, *R* = 2, *κ* = 0.1 and *λ* = 0.2.

In order to evaluate our ability to reconstruct transmission links we look at transmissions between observed hosts. Out of the 100 observed hosts, 67 are infected by another sampled individual. From our estimated transmission trees we consider both directional transmission links, where we must correctly establish the infector and infected host, and bidirectional transmission links, where a transmission link is established but the roles of infector and infected may swap. We define 0.5 as the posterior probability threshold for a transmission being identified, and define the sensitivity as the proportion of correctly identified transmission links (true positive rate). For the phylogenetic tree with one observation per host we obtain a sensitivity of 0.51 for bidirectional transmission links, and 0.28 for directional transmission links (Figure S1). For the phylogenetic tree with five observations per host the sensitivity increases to 0.64 for bidirectional transmission links, and 0.55 for directional transmission links (Figure 1). The specificity (true negative rate) is greater than 0.996 in all cases. The full distributions of posterior probability estimates in each setting are shown in Figure 2. Increasing the number of secondary observations allows us to better reconstruct transmission links, and crucially, to better distinguish the direction of transmission.

**Figure 1:**
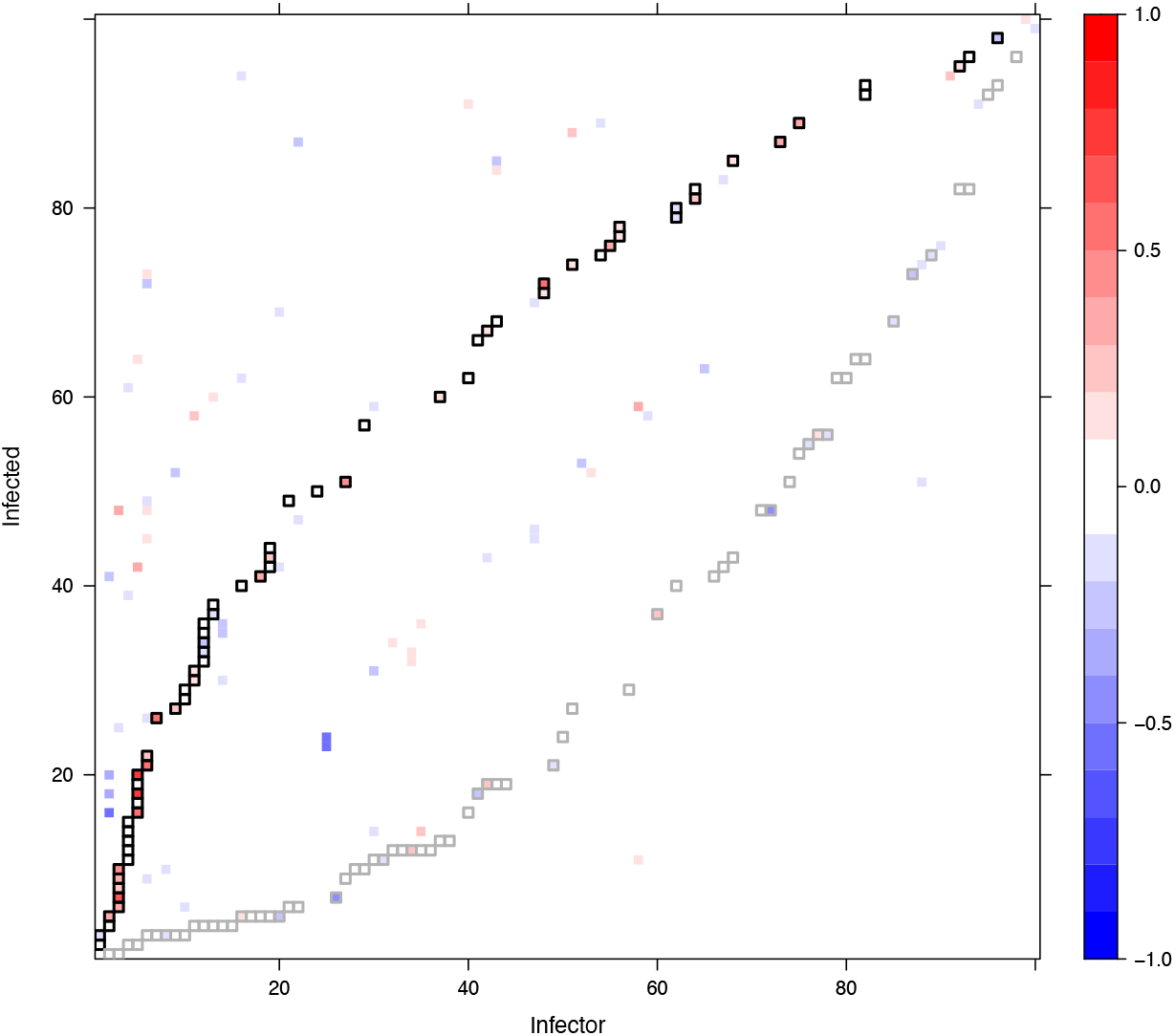
Difference in posterior probability estimates of transmission between a dataset with one observation per host and a dataset with five observations per host. The underlying transmission network remains the same; it is defined by the black squares, which show the true transmissions in the simulated dataset. The gray squares show the reverse relationship, switching the true infector and infected hosts. Black squares containing red demonstrate higher posterior probabilities being assigned to the true transmission links as a result of including more observations. Elsewhere, blue indicates lower posterior probabilities being assigned to incorrect transmission links.

**Figure 2:**
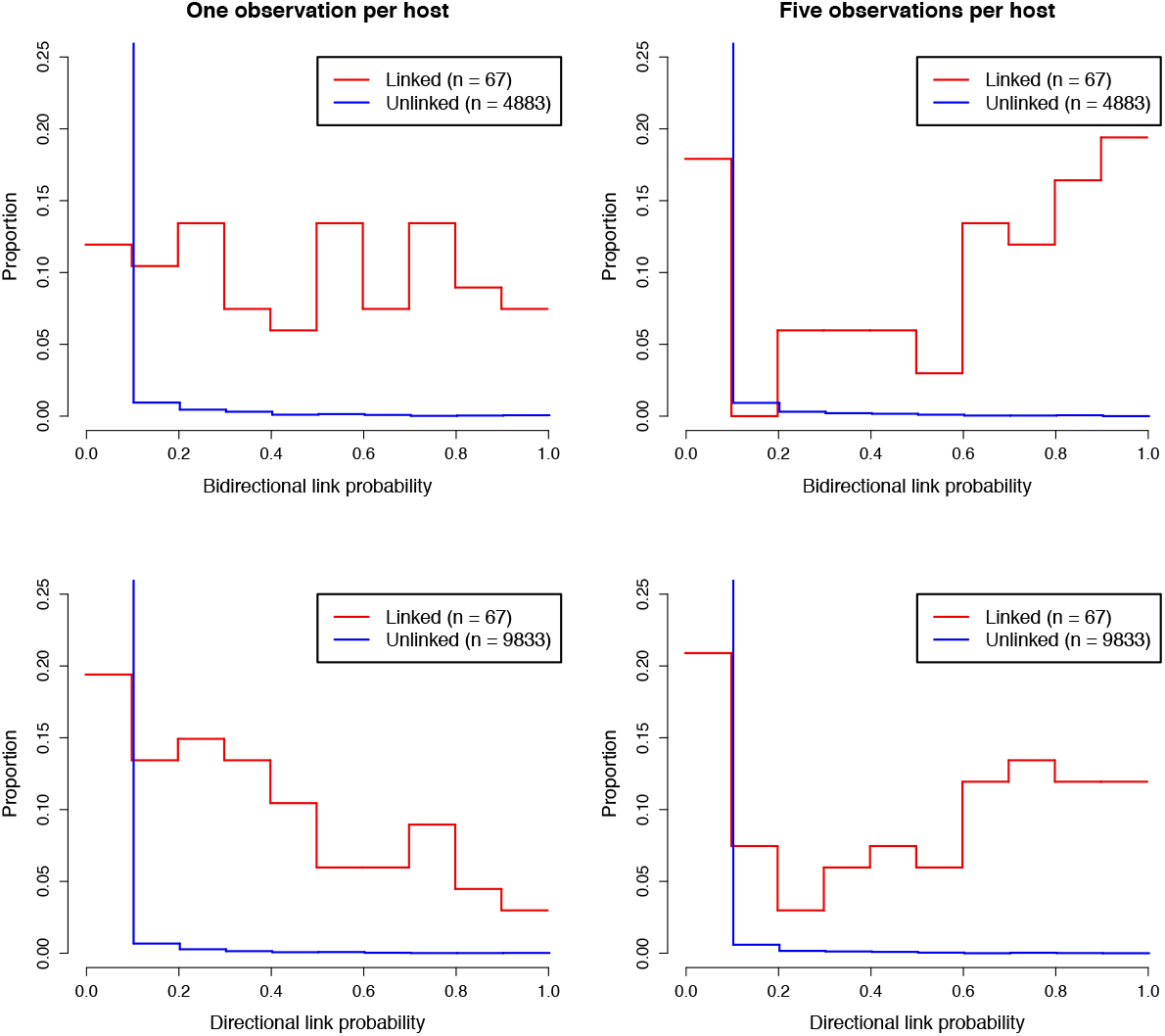
Distribution of posterior link probabilities inferred in the simulation studies with one (left) and five (right) observations per host. The top plots show bidirectional link probabilities in which the roles of infector and infected host may switch, the bottom plots show the directional link probabilities in which the infector and infected host must be correctly inferred. The red lines relate to pairs of individuals for which a transmission link exists, and the blue lines relate to pairs of individuals that are not linked.

The within-host population model plays a key role in our ability to establish transmission links. If the transmission of multiple lineages is more common, the posterior probabilities of transmission links will tend to be lower. For example, repeating the simulation process above with a full bottleneck (fixing *κ* = 0) results in a bidirectional (directional) sensitivity of 0.57 (0.43) with one observation per host, and 0.75 (0.63) with five observations per host, all higher than in the previous results with a partial bottleneck. On the other hand, increasing to *κ* = 0.4 leads to a bidirectional (directional) sensitivity of 0.34 (0.25) with one observation per host, and 0.54 (0.39) with five observations per host, all lower than the example with *κ* = 0.1.

### Benchmarking using multiple simulations

We now repeat this process, again using a simulated dataset with 100 hosts and five observations per host; but performing the inference on simulations generated from a range of key parameters (*π, R, λ*, and *κ*), totalling 43 datasets. As previously, both the generation time distribution and primary observation time distribution follow a Gamma distribution with shape parameter 2 and scale parameter 1, and secondary observations occur 0.25 years later than the previous sample.

For the MCMC chains we obtain 12,000 samples, and discard the first 2,000 as a burn-in. Figure 3 shows the posterior parameter estimates. The vertical lines show central 95% credible intervals for each parameter, and the posterior mean is shown with a solid circle. The horizontal and diagonal lines indicate the true parameter values used to generate the data. These results demonstrate strong performance of the algorithm across very different simulation settings.

**Figure 3:**
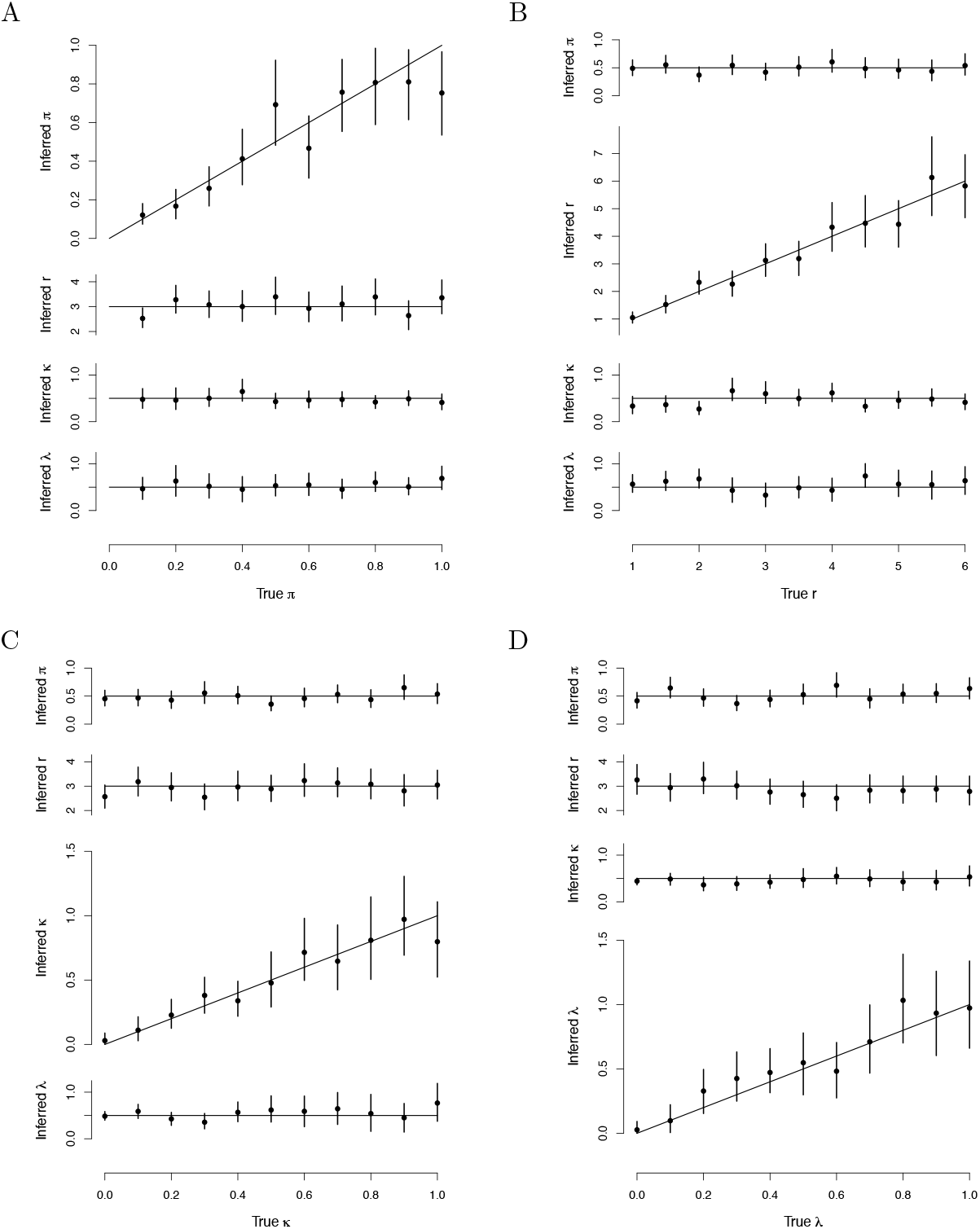
Varying the four key simulation parameters. Vertical bars show 95% central credible intervals, while solid circles show posterior means. Horizontal or diagonal lines show true values for simulations. (A) Varying *π*. (B) Varying *R*. (C) Varying *κ*. (D) Varying *λ*.

### Application to *Pseudomonas aeruginosa* transmission between cystic fibrosis patients

We reanalysed previously published genomic data from Danish cystic fibrosis (CF) patients infected with *Pseudomonas aeruginosa* (Marvig et al., 2013). This dataset included 42 genomes from 14 patients, sampled over almost 40 years between 1972 and 2008. Previous studies explored within-host evolutionary dynamics (Yang et al., 2011), variations in gene content (Rau et al., 2012) and comparative adaptation in CF human hosts (Marvig et al., 2013). The hosts are designated CFXXX as in these previous studies. We use as our starting point the dated phylogeny previously computed (Marvig et al., 2013) using BEAST (Suchard et al., 2018) and shown in Figure S2. It was previously noted (Yang et al., 2011) that one of the individual (CF66) had been infected twice in the 1970s and the 1990s, and so we modelled this as two separate hosts (labeled CF66a and CF66b). Infection with *P. aeruginosa* can be stable over long periods of time in CF patients (Rossi et al., 2021) and indeed some of the patients had been sampled, and found positive, over a period of more than 20 years (Marvig et al., 2013). We therefore set the generation time distribution to be Gamma with shape 2 and scale 5, resulting in a mean of 10 years, standard deviation of 7 years, and 95% range of 1.2 to 27.9 years. The last samples were from 2008 and the exact end of the sampling period was unclear from previous publications but we set it to the end of 2009.

We performed four separate runs of 100,000 iterations, which took approximately 3 hours on a standard laptop computer. For each of the four parameters *π, R, κ* and *λ* we checked that the effective sample size in each run was over 1,000 and the multivariate Gelman-Rubin statistic comparing runs was less than 1.1 (Brooks and Gelman, 1998). Figure 4A shows the dated tree, coloured by host according to the last state of one of the MCMC states. This colouring therefore represents a single draw from the posterior distribution of the transmission tree given the phylogeny. Changes in colours along the branches of the tree correspond to transmission events and are highlighted with red stars. Note that there are two simultaneous stars leading to the two genomes from patient CF180. These both correspond to infection from the same unsampled individual, since the yellow color found before the stars is not used by a leaf. Two lineages were transmitted from this donor to CF180 through the relaxed transmission bottleneck. Figure 4A is useful to illustrate the colouring process which relates the phylogenetic tree to the transmission tree. However, this only represents a single transmission configuration explored by the MCMC, and other iterations of the MCMC would look different, maybe with some of the same transmission events and others being different. It is therefore important to consider the probability of the transmission events. Figure 4B shows the matrix of probabilities of infection from each host to another, computed as the frequency of each transmission event across all MCMC iterations.

**Figure 4:**
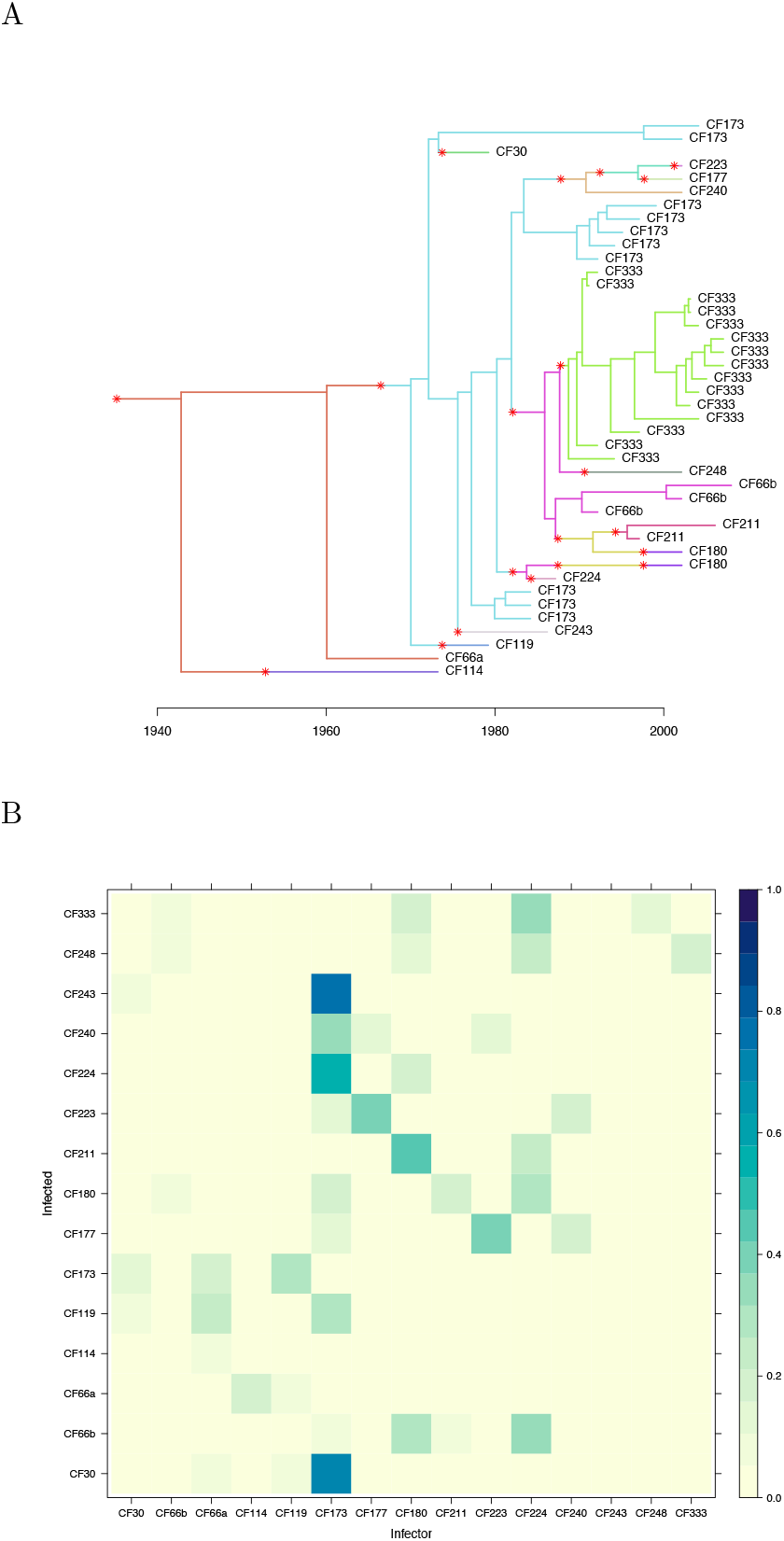
Transmission analysis of *P. aeruginosa*. (A) Dated phylogeny coloured by host according to a single draw from the posterior distribution. (B) Matrix of transmission probabilities from each host (row) to any other (column).

Figure S3 shows the trace and density of the parameters estimated in a single MCMC run. The sampling proportion was estimated to be *π* = 0.65, with a wide 95% credible interval [0.30 − 0.96]. The reproduction number was *R* = 1.20 [0.58 − 1.99]; as the credible interval includes one, it is not clear if the outbreak has the potential to cause a self-sustained epidemic. The within-host linear growth rate was *λ* = 0.56 [0.16 − 1.09] per year, which is lower than the prior exponential with mean one. On the other hand, the within-host starting population size was *κ* = 2.16 [0.41 − 5.05] which is higher than the prior exponential with mean one. This suggest that the bottleneck was not complete, and indeed attempting to fit the model with *κ* = 0 is impossible as it leads to a likelihood of zero. This is caused by the two samples from CF180 and the ten samples from CF173 being “inconsistent” as previously designated for samples from two hosts that cannot be explained by transmission of a single lineage (Romero-Severson et al., 2014, 2016). The individual CF173 was found to have infected at least three other hosts (CF30, CF224 and CF243) with probability higher than 50% (Figure 4B). These transmission events and their directionality are made clear by the paraphyletic relationship of the ten samples from CF173 as shown in Figure 4A (Leitner, 2019). In contrast, the 15 samples from CF333 formed a single monophyletic clade (Figure 4A) so that they are unlikely to have infected many others except maybe CF248 (Figure 4B).

### Application to a nosocomial outbreak of *Klebsiella pneumoniae*

An outbreak of carbapenem-resistant *Klebsiella pneumoniae* expressing the *bla*_OXA-232_ gene was identified over the course of 40 weeks at a single healthcare institution in California (Yang et al., 2017). A total of 17 infected patients were identified, from which 32 isolates were taken between 12th October 2014 and 17th July 2015. Case finding was performed using all samples in the 2014 and 2015 calendar years (Yang et al., 2017) and so we set the date for the end of the sampling period to the end of 2015. Whole-genome sequencing was applied to these *K. pneumoniae* isolates and a dated phylogeny was computed previously (Yang et al., 2017) using BEAST (Suchard et al., 2018) which is shown in Figure S4. The hosts are labeled either PtXXX if they were symptomatic or CPtXXX if they were colonized, as in the previous study (Yang et al., 2017). We set the generation time distribution to be exponential with mean 0.5 year, following a previous study of another *K. pneumoniae* hospital outbreak (van Dorp et al., 2019). We used the same number of MCMC runs, length of runs, and convergence diagnostics as in the previous application.

Figure S5 shows the trace and density of the parameters estimated in a single MCMC run. The sampling proportion was estimated to be high, with *π* = 0.88 [0.60 − 0.99], suggesting that there were only few missing transmission links between the 17 sampled patients. The basic reproduction number was *R* = 0.97 [0.37 − 1.74], with the credible interval including the value of one needed for an outbreak to spread beyond a few cases. The within-host linear growth rate was *λ* = 0.49 [0.03 − 1.28] per year and the within-host population size at time of infection was *κ* = 0.066 [0.009 − 0.158]. This is lower that the prior exponential with mean one and suggests that the transmission bottleneck was almost complete during this small outbreak. However, the transmission bottleneck was not absolutely complete, as indicated by the fact that fitting our model with *κ* = 0 would result in a likelihood equal to zero. This is because the six samples from Pt6 and the two samples from Pt9 are inconsistent, as can be seen in the dated phylogeny on Figure S4.

Figure 5A shows the dated tree coloured by host at the last MCMC iteration, while Figure 5B shows the posterior probabilities of infection from any host to any other. Strikingly, according to our analysis patient Pt6 had a greater than 50% posterior probability of having infected seven other patients (CPt2, CPt4, CPt5, CPt6, Pt5, Pt7 and Pt9). There were six genomes isolated from Pt6, with dates ranging from 7th January 2015 to 17th July 2015 which is more than half of the overall sampling period. The specimen types for these isolates were quite diverse: three from blood, one rectal and two from bile (Yang et al., 2017), suggesting that the patient was infected long enough for the pathogen to spread throughout their body. While other patients in the study do present a similar number of samples, a comparable variety of originating tissues, and a similarly long infection duration — for instance patient Pt1, with seven genomes from respiratory, abdominal and blood specimen over a period of several months — that does not translate in a similar amount of infection events estimated by our method. In fact, the genetic diversity of isolates from Pt6 appears to be very high (Figure 5A), thus backing our inference that Pt6 is a superspreading individual (Lloyd-Smith et al., 2005). This could not have been detected without the use of multiple genomes.

**Figure 5:**
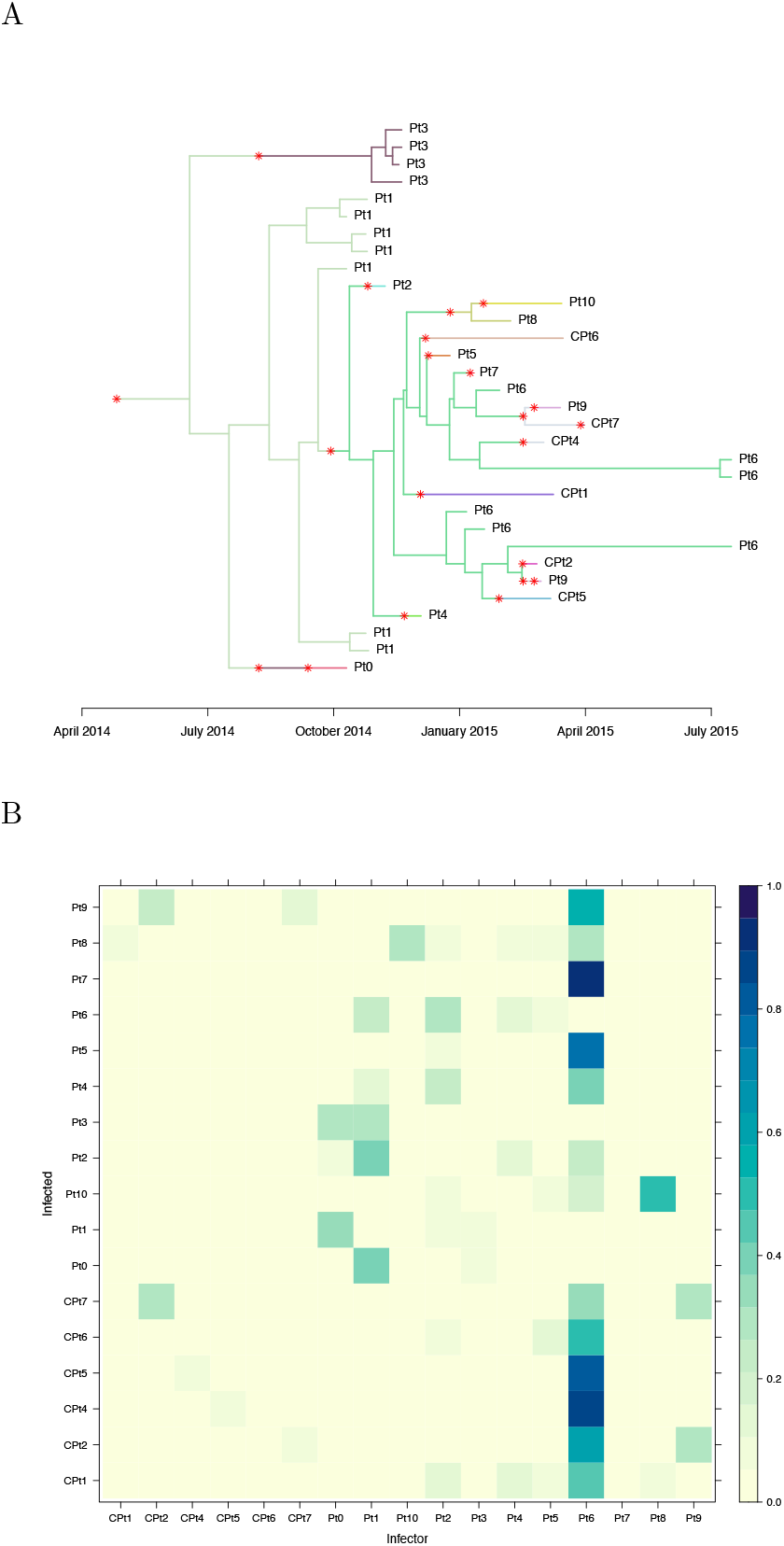
Transmission analysis of *K. pneumoniae*. (A) Dated phylogeny coloured by host according to a single draw from the posterior distribution. (B) Matrix of transmission probabilities from each host (row) to any other (column).

## DISCUSSION

We have described new methodology for inferring who infected whom from a dated phylogenetic tree in which hosts have potentially been sampled multiple times. A key change compared to previous work (Didelot et al., 2014, 2017) is the removal of the full transmission bottleneck, meaning that hosts may be infected with multiple lineages from the transmission donor. Without this change many phylogenetic trees with multiple samples per host would not support compatible transmission trees (Romero-Severson et al., 2014, 2016; Leitner, 2019). Indeed the two real datasets we analysed, corresponding to outbreaks of *Pseudomonas aeruginosa* and *Klebsiella pneumoniae*, could not be explained without relaxing the transmission bottleneck. Most previous transmission analysis methods could not accommodate more than a single genome per host, so that leaves would need to be pruned from the phylogenetic tree in order to undertake transmission inference (Xu et al., 2020), leading to less informative outcomes. Under our new methodology we are able to incorporate multiple samples per host, resulting in the stronger identification of transmission links and their direction, as was showed when analysing simulated datasets.

A few methods have been proposed recently to exploit information about within-host diversity contained in deep sequencing data of a single sample (De Maio et al., 2018; Wymant et al., 2018; Ortiz et al., 2022), but this is quite different and potentially less informative than using genomes from multiple samples. The only available tool we are aware of that can use multiple genomes to inform a transmission analysis is SCOTTI (De Maio et al., 2016). However, this is based on a very different approach to the one we took, namely using an approximation to the structured coalescent (Muller et al., 2017) called BASTA which was originally developed for phylogeographic reconstruction (De Maio et al., 2015). Here instead we build upon previous work (Didelot et al., 2014, 2016) that performs transmission analysis by colouring the branches of a pre-established dated phylogeny. This allows us to model the relationship between transmission tree and phylogeny through an explicit within-host evolutionary model, to develop an explicit transmission model in which sampled and unsampled individuals are featured, and to achieve better scalability by separating phylogenetic inference from its epidemiological interpretation (Didelot and Parkhill, 2022).

Our method implements a general pathogen population growth model rather than using the constant bounded coalescent model (Carson et al., 2022). The main restriction on the choice of model is that we must be able to calculate the likelihood of the phylogenetic tree, which in turn means that the coalescence rate must be integrable. However, this is not a strong requirement, as many widely used models satisfy it — among them the exponential growth model, the logistic growth model, or any piecewise models with separate growth and decay phases. For the work presented here we used a linear growth model, which has been used before in HIV work (Romero-Severson et al., 2014, 2016; Leitner, 2019), but for most other pathogens there is little information about which within-host population size model is most realistic (Didelot et al., 2016). We demonstrated that using phylogenetic trees with multiple samples per host improves the estimation of the population model parameters. With sufficient samples per host it should be possible to determine which within-host population size models are more strongly supported by the data, for example and comparing the evidence of each model (Friel and Wyse, 2012).

Our methodology maintains some of the assumptions from previous work (Didelot et al., 2017), for example the sampling proportion and reproduction number are assumed to remain constant through time. In many settings, users would have knowledge about whether and how the sampling proportion varied over time, for example by looking at the number cases for which genomic sequences are available divided by the number of confirmed cases (Jelley et al., 2022). This information could be integrated relatively easily into an analysis, by having users supply a function *π*(*t*) instead of the constant *π*. On the other hand, it would often be interesting to infer variations in the reproduction number *R*(*t*), since this would provide an additional genomic-based estimate compared to existing methods based on incidence data (Wallinga and Teunis, 2004; Cori et al., 2013). A simple approach would be to use a step-wise constant function. The dates of these steps may be fixed based on real-world policy changes, such as intensifying monitoring in response to an outbreak, or potentially inferred via change point detection (Tartakovsky and Moustakides, 2010).

In conclusion, we presented a new Bayesian inference method for the reconstruction of transmission trees from dated phylogenetic trees in which hosts are sampled multiple times. This method is implemented in a R package that extends TransPhylo and is available at https://github.com/DrJCarson/TransPhyloMulti. When applied to longitudinally sampled genomes from several infected individuals, our method has the potential to improve our understanding of both the within-host and between-host dynamics of many pathogens causing infectious disease.

## MATERIALS AND METHODS

### Notation

Let us denote 𝒫 as the dated phylogenetic tree, 𝒯 as a transmission tree, *θ*_*P*_ as the coalescent model parameters, and *θ*_*T*_ as the transmission model parameters. We want to sample from the posterior distribution

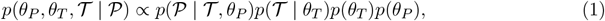

where the term *p*(𝒫 | 𝒯, *θ*_*P*_) is the likelihood of the coalescent model conditional on a given transmission tree, the term *p*(𝒯 | *θ*_*T*_) is the likelihood of the transmission model, and the terms *p*(*θ*_*P*_) and *p*(*θ*_*T*_) are prior distributions.

We parameterise the transmission tree 𝒯 as follows. Let *x* be a vector of infection times such that element *x*^*j*^ gives the infection time of host *j*. Likewise let *A* be a vector of infectors, so that if *A*^*j*^ = *i* then host *j* was infected by host *i*. We indicate the root host by setting *A*^*j*^ = 0. Primary observation times are denoted by vector *y*, with the corresponding host denoted by vector *H*_*y*_. Secondary observation times are denoted by vector *z*, with host *H*_*z*_.

For the phylogenetic tree 𝒫 we need to consider the leaf and coalescent times. The leaves correspond to observations under the transmission tree. We denote the vector of leaf times *s* and corresponding hosts *H*_*s*_, noting that *s* = (*y, z*) and that *H*_*s*_ = (*H*_*y*_, *H*_*z*_). We indicate the coalescent ancestor using vector *C*_*s*_. The coalescent times are denoted by vector *u*, and parents *C*_*u*_. We again denote the root node with 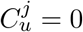.

Figure 6A demonstrates a transmission tree with

**Figure 6:**
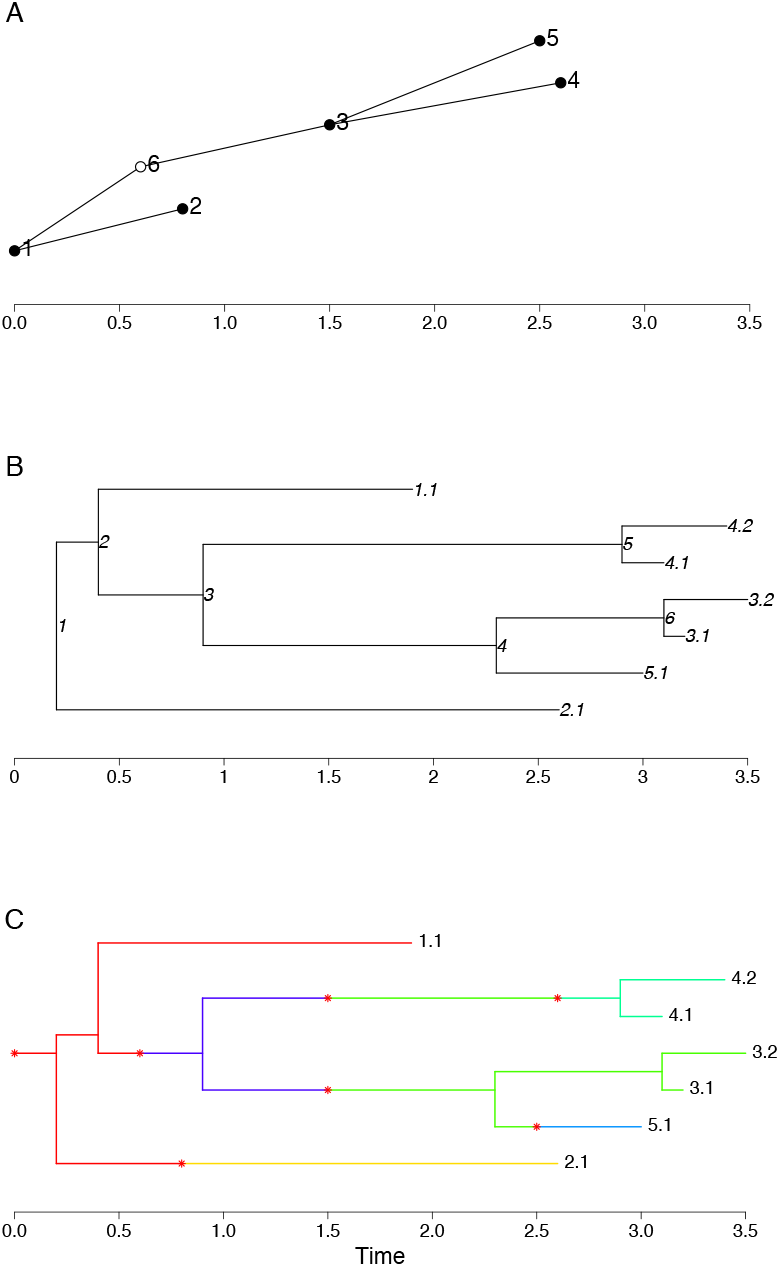
(A) Example transmission tree with six hosts. Points indicate the infected times of each host. Filled circles show observed hosts, and empty circles show unobserved hosts. (B) Example phylogenetic tree with seven leaves from five observed hosts. Leaf labels indicate the host, followed by the sample number for that host. Each coalescence node is given a label. (C) Example coloured phylogenetic host with seven leaves from five observed hosts, and six hosts overall. The branch colour indicates the host, and the asterisks indicate transmissions. Here host 3 is infected with two lineages.

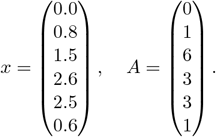

That is, host 1 infects hosts 2 and 6, host 6 infects host 3, and host 3 infects hosts 4 and 5. In addition we have primary and secondary observations (not shown), for example

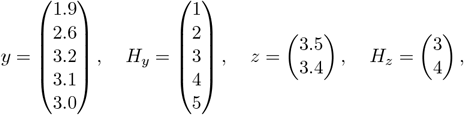

indicates that hosts 1, 2 and 5 are observed once, hosts 3 and 4 are observed twice, and host 6 is unobserved.

Figure 6B shows an example phylogenetic tree obtained by combining the primary and secondary

observations from the transmission tree. Here

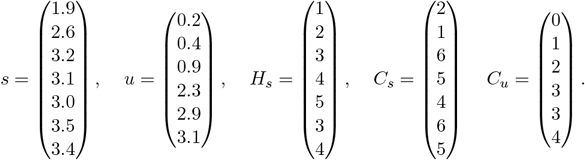

We can represent both the transmission and phylogenetic trees as a coloured phylogenetic tree, as shown in Figure 6C. Doing so highlights that each coalescent event is now assigned to a host. We denote these hosts as *H*_*u*_, in this example

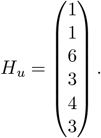

### Epidemiological model

The epidemiological model is a stochastic branching process in which infected individuals transmit to secondary cases (offspring). The number of offspring *k* is sampled from the offspring distribution *α*(*k*), assumed to be a negative binomial distribution with parameters (*r, p*), i.e.

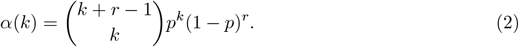

The time between the primary and any secondary infection is sampled from the generation time distribution *γ*(*τ*), which typically follows a Gamma distribution with known parameters.

Under a *finished outbreak* scenario, each host is assumed to be observed with probability *π*. The time between the host being infected and first being observed is sampled from the observation time distribution *σ*(*τ*). As with the generation time distribution this is typically a Gamma distribution with known parameters.

In some applications observations occur over a restricted time interval, or possibly set of time intervals. In such applications the probability of a host being observed depends on their infection time. An example we will look at is the *ongoing outbreak* scenario, in which there is an observation cut-off time *T*. In this scenario a host infected at time *t* is observed with probability

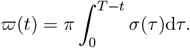

In other words, we use the same observation distribution as the finished outbreak scenario, but treat observations later than *T* as censored.

Finally, hosts may be observed multiple times. We assume that any host can only be infected once, and that any subsequent observations relate to the same infectious period. We define *β*(*b*) as the distribution for the number of secondary observations *b* ≥ 0, and *ρ*(*τ*_1:*b*_) as the distribution for the times between the secondary observations and the primary observation assuming that *b* ≥ 1. This is an additional modelling component to the previous version of TransPhylo (Didelot et al., 2017). However, by assuming that the secondary observation times depend only on the primary observation times, we can undertake inference in a similar manner without formally specifying these distributions. Under our modelling assumptions we can express the likelihood of the transmission tree as

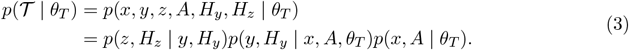

Consequently, the term *p*(*z, H*_*z*_ | *y, H*_*y*_) will cancel in any inference scheme exploring the parameter space (*x, A, θ*_*T*_), allowing us to drop this element of the transmission tree likelihood.

### Host inclusion and exclusion

Our goal is to infer a transmission tree from a dated phylogenetic tree. This can be visualised as *colouring* the branches of the phylogenetic tree, where each colour represents a distinct host. For a host to appear on the phylogenetic tree they must either be observed directly or be an ancestor to a different observed host. We refer to such hosts as *included* hosts. In many applications the number included hosts is dwarfed by the number of hosts implied by the epidemiological model to not appear on the phylogenetic tree (*excluded* hosts). Examples include when *π* is small, or when *r* is large in an ongoing outbreak scenario. In the latter case, a large number of hosts will be infected shortly before the observation cut-off time, and so will be excluded with high probability. For this reason we instead formalise a transmission model for only the included hosts.

Define *ω*(*t*) as the exclusion probability of a host infected at time *t*. Assuming that *T* is the cut-off time for observations *ω*(*t*) = 1 for *t* ≥ *T* . We can then define the following recursive relationships.

The exclusion probability of an offspring from a host infected at time *t* is

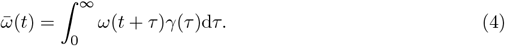

The probability that all offspring from an individual infected at time *t* are excluded is

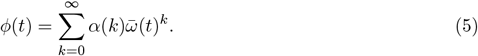

The exclusion probability of an individual infected at time *t* is

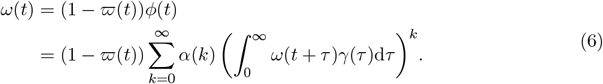

That is, the probability of the host being unobserved and having no included offspring. In the finished outbreak scenario the recursive relationship is simply

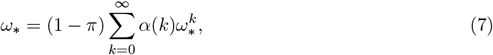

with *ω*_*_ being the exclusion probability for every host. Note that these calculations do not depend on the secondary observation times or their distribution.

### Numerical approximations

The exclusion probabilities are intractable, and so we use numerical approximations. For example, consider the ongoing outbreak scenario with observation cut-off time *T* . For *t* ≥ *T, ω*_*t*_ = 1, and so

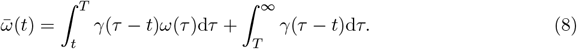

The second term can be computed explicitly, and the first term can be approximated using the trapezoid method:

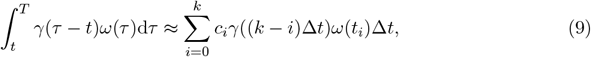

where *c*_*i*_ = 1 for 0 *<i<k* and *c*_*i*_ = 0.5 otherwise, and *t*_*i*_ = *T* − *i*Δ*t*. Assuming *γ*(0) = 0:

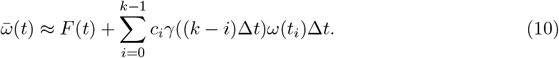

where 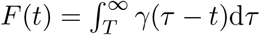.

Using the probability generating function of a negative binomial distribution with parameters *r* and *p*, we can evaluate

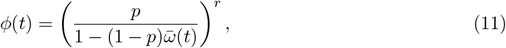

and finally

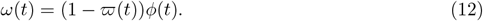

Both will be approximate owing to the approximation of 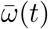. All three exclusion probabilities are therefore approximated by iterating backwards through time from *T* in discrete steps of size Δ*t*.

### Transmission tree likelihood

We can now define a likelihood for the transmission tree for only included individuals. Throughout we will set *T* as the cut-off time for observations. Consider first the root case (the first infected individual in our transmission chain) with infection time *x*^1^, and let *I*^1^ = 1 denote that the root case is included. The probability that the root host is unobserved (denoted by *S*^1^ = 0) given that they are included is

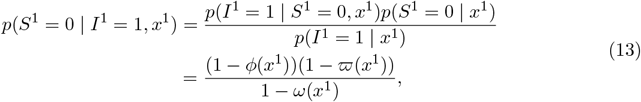

and the probability that the root case is observed (*S*^1^ = 1) is

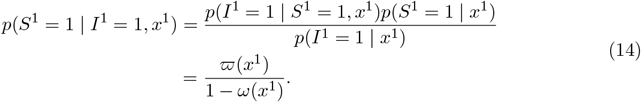

In the event the root case is observed we also need to calculate the density of the primary observation time *y*^1^,

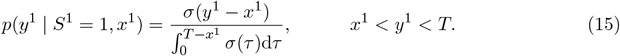

Additionally the full transmission tree likelihood incorporates the density of the secondary observation times. However, when it comes to undertaking inference these terms will cancel out, and so we skip this step.

Second, we calculate the probability that the root case has *d*^1^ included offspring. The probability of a host infected at time *t* producing *d* included offspring is

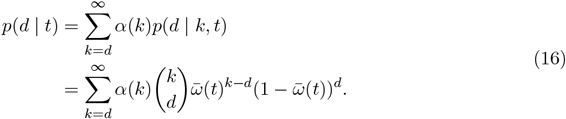

We then need to condition on whether or not the root case was sampled. If the root case was not sampled, they must produce at least one included offspring to be included, and so

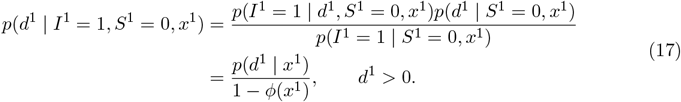

If the root case was sampled, then it is included for any value of *d*^1^, and so

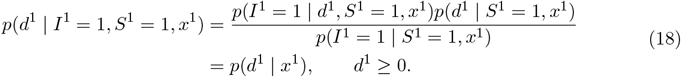

In the event *d*^1^ *>* 0, we also calculate the density of the transmission times for any included offspring. Denoting ℋ^1^ as the offspring labels, 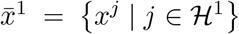 as the set of offspring infection times, and *Ī*^1^ = 1 that the set of offspring are included, the likelihood contribution is

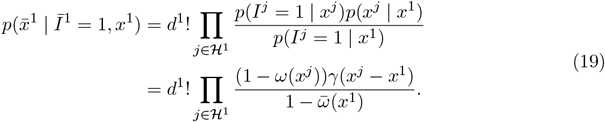

The *d*^1^! term arises from the fact that the ordering of the offspring infection times is inconsequential.

In summation, the likelihood contribution (sans secondary observations) for the root host in the unobserved case is

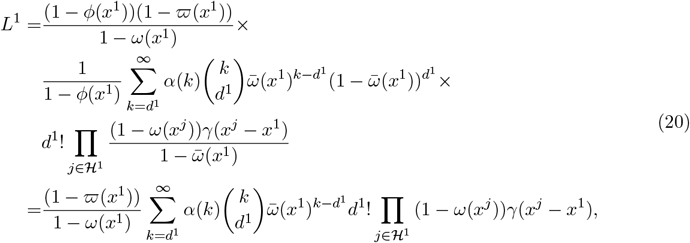

and for the observed case is

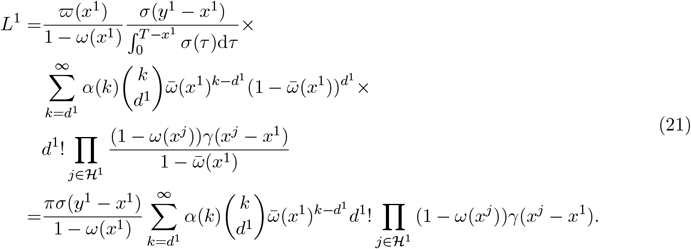

The full likelihood is calculated by induction, applying the same density calculations to each included host. Note that in doing so, with the exception of the root host, the terms 1 − *ω*(*x*^*j*^) will cancel in the likelihood. The likelihood of the finished outbreak scenario can be calculated using a similar approach.

### Transmission tree simulation

There are multiple ways to simulate transmission trees in the ongoing outbreak scenario. One option is to fix the observation cut-off time *T* and follow the steps used in the likelihood derivation in Equations (13) to (19). This is presented in Algorithm S1. Alternatively, an upper bound may be set for either the number of observed hosts or the number of observations, with *T* being a simulated outcome. The transmission tree is simulated from the stochastic branching model, from the earliest infected host to latest, until this relevant bound is exceeded.

This induces a temporary cut-off time *T*′. Simulation then continues so long as there are hosts infected earlier than *T*′ for which the number of observations and number of offspring have not been generated. Finally, a suitable value for *T* is generated that returns the correct number of observed hosts / observations. When bounding by the number of observed hosts *T* acts as a cut-off time for primary observations only, whereas when bounding by the number of observations *T* acts as a cut-off time for both primary and secondary observations. Further details are given in Algorithms S2 and S3.

### Coalescent model

In the original version of TransPhylo the coalescent model used was the bounded coalescent (Carson et al., 2022). This model follows the standard coalescent model with heterochronous sampling (Drummond et al., 2002), but conditions all lineages to coalesce before the infection time of each host. Here we need to choose a coalescent model that allows for the transmission of multiple lineages between hosts. With a bottleneck assumption many dated phylogenetic trees would not permit the overlaying of a transmission tree under our stochastic branching model.

Here we assume that the within-host pathogen population size *q*(*τ*) grows linearly:

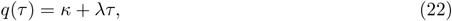

where *τ* is the time since the host was infected. Should *κ* = 0 all lineages will coalesce by the host’s infection time. We could adopt alternative population models, so long as they are integrable.

### Phylogenetic tree likelihood

The likelihood of the phylogenetic tree conditional on the set of transmissions is calculated by taking the product of the likelihood of each *subtree* for each host. The subtree of any host *j*, denoted here as 𝒫_*j*_, is formed by taking the parts of the phylogenetic tree assigned (coloured) by host *j*. Each subtree is rooted at the host’s infection time *x*^*j*^, with the number of roots being the number of lineages transmitted to the host. Leaves correspond to observations of the host and transmissions to the hosts included offspring, noting that each transmission may contribute multiple leaves (transmitting multiple lineages).

Let 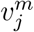, *m* = 1, …, *M*_*j*_ be the times leaves are added within the subtree of host *j*, and let 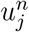, *n* = 1, …, *N*_*j*_ be the coalescence times, supposing *N*_*j*_ *>* 0. Then we define the number of extant lineages at time *t* as4

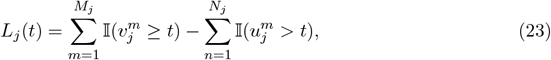

so that if *t* is the time of a coalescence, *L*_*j*_(*t*) is the number of lineages that could have coalesced.

Denoting *τ*_*j*_ = *t* − *x*^*j*^, the likelihood contribution from each host is then

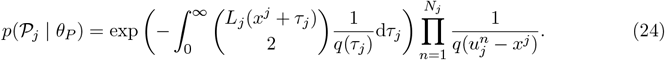

Let 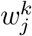, *k* = 0, …, *K* be the ordered set of root, leaf, and coalescence times, with 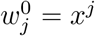. Let 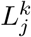 be the number of lineages in the interval 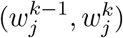. The integral in the exponent can then be partitioned accordingly

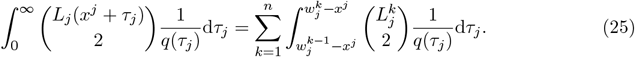

For the linear growth model, these terms are then

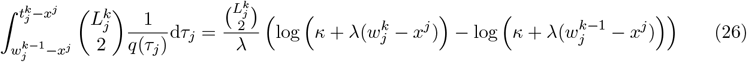

### Phylogenetic tree simulation

The subtree for each host is simulated using repeated application of inverse transform sampling. Assume that at time *t* there are *L* extant lineages in a host infected at time *x*^*j*^. We wish to sample the next coalescence time 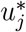, and so we solve

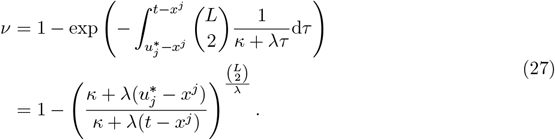

Thus, we sample *V* ∼ 𝒰(0, 1) and propose

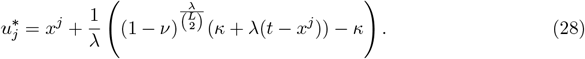

Should additional leaves be added at time 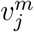, where 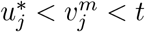, then the proposed coalescence time is discarded, and the next coalescence time is proposed by setting 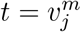 and *L* = *L* + 1.

If 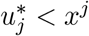 then the coalescence time is discarded, and the *L* lineages are assumed to have been transmitted to the host. Otherwise, we accept the proposed coalescence time and randomly select a pair of lineages to coalesce. We then propose the next coalescence time by setting 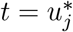 and *L* = *L* − 1. The complete algorithm is presented in Algorithm S4.

### Inference

Inference is undertaken using reversible-jump Markov chain Monte Carlo (Green, 1995). We iterate through the following update steps:

1. Update the transmission model parameters according to *p*(*θ*_*T*_ |𝒯).
2. Update the coalescent model parameters according to *p*(*θ*_*P*_ | 𝒫).
3. Update the transmission tree according to *p*(𝒯 | 𝒫, *θ*_*T*_, *θ*_*P*_).

Steps 1 and 2 are performed using multivariate Gaussian random walks, conditional on the current transmission and phylogenetic trees. The scale and covariance in each case is determined using the accelerated shaping and scaling algorithm of Spencer (2021) with target acceptance *a* = 0.234 and forgetting sequence *f* (*n*) = l0.5*n*j.

In Step 3 we randomly select from three proposals that update the transmission tree conditional on the current model parameters: an add proposal for adding a new transmission to the current transmission tree, a remove proposal for removing a transmission, and a local move proposal that updates the infection time of a single host and potentially exchanges offspring with its infector. Each proposal ensures that the new transmission tree is consistent with the phylogenetic tree. Further details are provided below.

Step 3 makes relatively small changes to the transmission tree with each update. Additionally, the computational cost is relatively cheap as we only need to evaluate the likelihood contributions from the one or two affected hosts. Consequently it is beneficial to perform Step 3 multiple times in each scan, in order to improve the mixing of the MCMC. In general, we find that performing O(*N*) Step 3 updates in each scan works well, where *N* is the number of primary observations.

### Add proposal

In the add proposal a new transmission is added to the transmission tree. We can interpret this proposal as sampling the following components:

1. Sampling the infector of the new host from the current hosts.
2. Sampling which of the infector’s offspring become the offspring of the new host.
3. If the infector is observed under the current transmission tree, determine whether those observations are assigned to the infector or the new host.
4. Sampling a transmission time (the infection time of the new host).

Note that the new host must be included, requiring at least one offspring or observation. Additionally, the phylogenetic tree constrains which set of offspring can be infected by the new host, and the set of times over which the transmission can occur. The combined sampling information also determines how coalescent events are divided between the two hosts.

Firstly, we sample the infector uniformly from the set of current hosts. We then consider how a transmission event can be added to the infector’s subtree. The division of the offspring and observations between the two hosts is determined by the branch or branches over which the transmission occurs. Consequently, the discrete parts of our proposal corresponds to sampling this set of branches.

Let Ω denote the set of all possible combinations of branches of the subtree, such that |Ω| = 2^*B*^ − 1 where *B* is the total number of branches. Let *V* ^*j*^ indicate the length of overlap for combination *j* ∈ {1, …, 2^*B*^ ‒ 1}. A particular combination is then sampled according to the probabilities *V*^*j*^/∑_*j*_ *V*^*j*^. Whilst this seems like an expensive calculation, note that most combinations have no overlap. Additionally, some branches can be pre-grouped. For instance, observations cannot be split between hosts, so transmissions would be required to occur on all branches leading to an observation, or none.

Once a combination of branches has been sampled, let 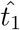 denote the earliest time of overlap of the group, and 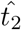 the latest. The transmission time is then sampled uniformly between these values.

As an example, consider adding a transmission to the coloured phylogenetic tree depicted in Figure 7A with Host 1 as the infector. We will refer to the newly added host as Host P. There are three branches within the subtree and one possible combination of two branches with non-zero overlap. This gives us four possible placements of the new transmission:

**Figure 7:**
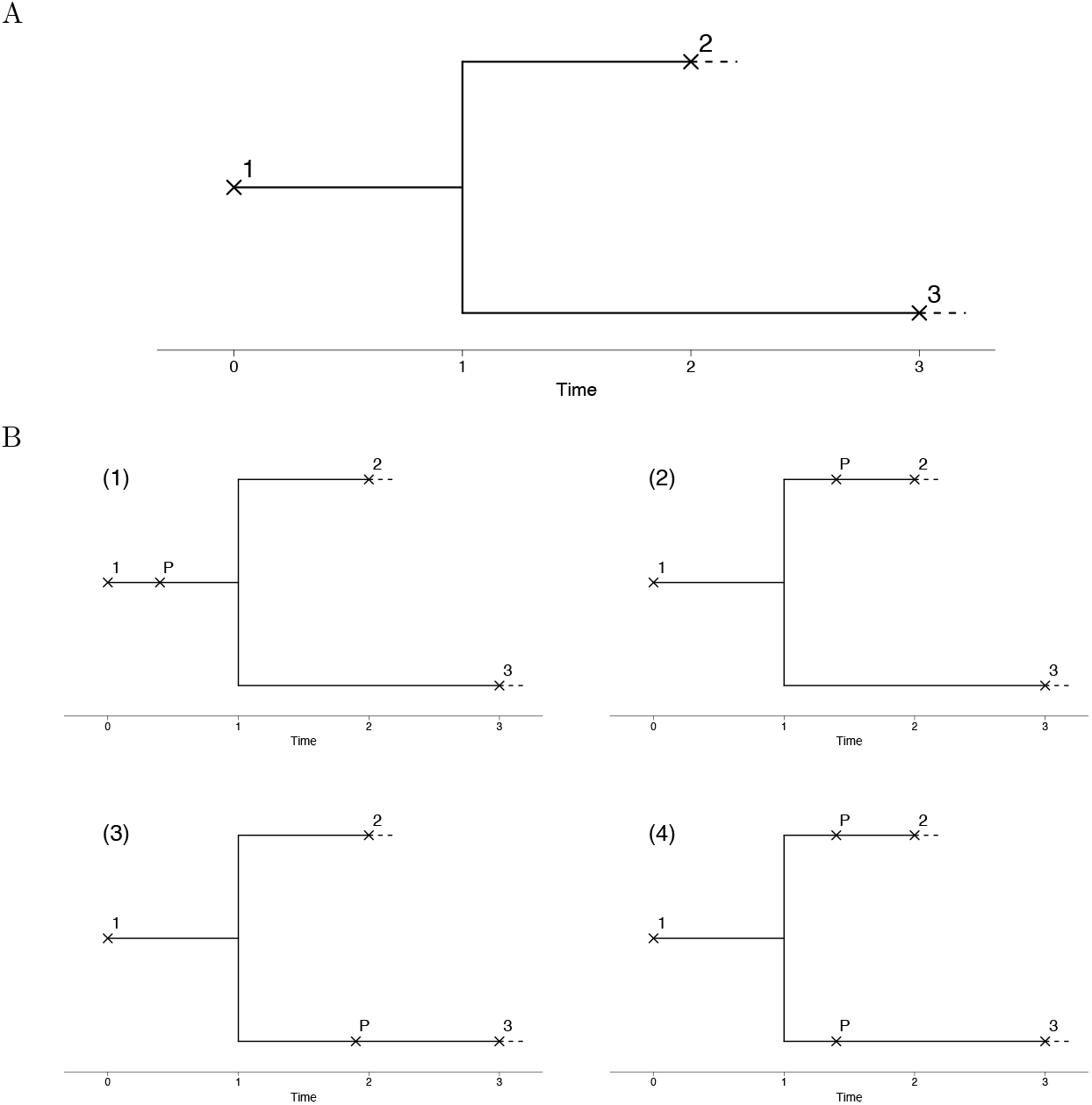
(A) Part of a coloured phylogenetic tree focused on host 1. Transmissions are indicated by × and labelled by the infected host number. (B) Examples of the three topologies when adding a transmission. Transmissions are indicated by × and labelled by the infected host number. Under the existing transmission tree, Host 1 infects both Host 2 and Host 3. When adding a transmission, another host is added, denoted as Host P.

1. Between the infection time of Host 1 and the coalescent, on the branch leading to the coalescent.
2. Between the coalescent and the infection time of Host 2, on the branch leading to the infection of Host 2.
3. Between the coalescent and the infection time of Host 3, on the branch leading to the infection of Host 3.
4. Between the coalescent and the infection time of Host 2, on the two branches leading to the infections of Host 2 and Host 3.

These different options are shown in Figure 7B. Note that placement 1 and 4 describe the same transmission tree. Host 1 infects Host P, and Host P infects Hosts 2 and Host 3. However, the number of transmitted lineages, and in which host the coalescent lies, depends on the exact placement. Evaluating *V* ^1^ = 1, *V* ^2^ = 1, *V* ^3^ = 2, *V* ^4^ = 1 gives us the probability vector (0.2, 0.2, 0.4, 0.2) for sampling the placement.

### Remove proposal

In the remove proposal we first determine which transmissions can be removed such that the new transmission tree remains consistent with the phylogenetic tree. In particular this means that we can not remove transmissions between two observed hosts, since these observations would then be assigned to a single host. From this subset, and excluding the root transmission, a single transmission is sampled at random and removed.

### Local move (remove-add) proposal

Under the local move proposal we aim to move a transmission with minimal changes to the transmission tree topology. This can also be seen as a remove-add proposal, where we first remove a transmission, and then add a new transmission nearby. The remove step is similar to the remove proposal, but any transmission can be sampled. We are no longer concerned about whether removing a transmission results in an inconsistent transmission tree, as we can ensure consistency when the transmission is re-added. If the root transmission is sampled, a new root time is proposed using a Gaussian random walk. Otherwise, the selected transmission is removed, and a new transmission added using the add proposal with the same infector.

### Implementation

We implemented the methods above into a new R package called TransPhyloMulti which extends TransPhylo. TransPhyloMulti is available at https://github.com/DrJCarson/TransPhyloMulti. This repository also contains all the code and data needed to reproduce all results shown in this paper. The R package ape was used to store, manipulate and visualise phylogenetic trees (Paradis and Schliep, 2019).

## ACKNOWLEDGEMENTS

We acknowledge funding from the National Institute for Health Research (NIHR) Health Protection Research Unit in Genomics and Enabling Data (grant number NIHR200892).

## References

Biek R, Pybus OG, Lloyd-Smith JO, Didelot X. 2015. Measurably evolving pathogens in the genomic era. Trends in Ecology & Evolution. 30:306–313.

Boeras DI, Hraber PT, Hurlston M, Evans-Strickfaden T, Bhattacharya T, Giorgi EE, Mulenga J, Karita E, Korber BT, Allen S, et al. (11 co-authors). 2011. Role of donor genital tract HIV-1 diversity in the transmission bottleneck. Proceedings of the National Academy of Sciences. 108:E1156–E1163.

Bouckaert R, Vaughan TG, Barido-Sottani J, Duchêne S, Fourment M, Gavryushkina A, Heled J, Jones G, Kühnert D, De Maio N, et al. (25 co-authors). 2019. BEAST 2.5 : An Advanced Software Platform for Bayesian Evolutionary Analysis. PLoS Computational Biology. 15:e1006650.

Brooks SP, Gelman A. 1998. General methods for monitoring convergence of iterative simulations. Journal of Computational and Graphical Statistics. 7:434–455.

Bryant JM, Grogono DM, Greaves D, Foweraker J, Roddick I, Inns T, Reacher M, Haworth CS, Curran MD, Harris SR, et al. (13 co-authors). 2013. Whole-genome sequencing to identify transmission of Mycobacterium abscessus between patients with cystic fibrosis: A retrospective cohort study. The Lancet. 381:1551–1560.

Campbell F, Didelot X, Fitzjohn R, Ferguson N, Cori A, Jombart T. 2018. Outbreaker2: A Modular Platform for Outbreak Reconstruction. BMC Bioinformatics. 19:363.

Carson J, Ledda A, Ferretti L, Keeling M, Didelot X. 2022. The bounded coalescent model: Conditioning a genealogy on a minimum root date. Journal of Theoretical Biology. 548:111186.

Cori A, Ferguson NM, Fraser C, Cauchemez S. 2013. A new framework and software to estimate time-varying reproduction numbers during epidemics. American journal of epidemiology. 178:1505–12.

Cortey M, Ferretti L, Pérez-Martín E, Zhang F, de Klerk-Lorist LM, Scott K, Freimanis G, Seago J, Ribeca P, van Schalkwyk L, et al. (11 co-authors). 2019. Persistent infection of African buffalo (Syncerus caffer) with foot-and-mouth disease virus: limited viral evolution and no evidence of antibody neutralization escape. Journal of virology. 93:10–1128.

Cottam EM, Wadsworth J, Shaw AE, Rowlands RJ, Goatley L, Maan S, Maan NS, Mertens PPC, Ebert K, Li Y, et al. (18 co-authors). 2008. Transmission pathways of foot-and-mouth disease virus in the United Kingdom in 2007. PLoS Pathogens. 4:e1000050.

De Maio N, Worby CJ, Wilson DJ, Stoesser N. 2018. Bayesian reconstruction of transmission within outbreaks using genomic variants. PLOS Computational Biology. 14:e1006117.

De Maio N, Wu CH, O’Reilly KM, Wilson D. 2015. New Routes to Phylogeography: A Bayesian Structured Coalescent Approximation. PLoS Genetics. 11:e1005421.

De Maio N, Wu CH, Wilson DJ. 2016. SCOTTI: Efficient Reconstruction of Transmission within Outbreaks with the Structured Coalescent. PLoS Computational Biology. 12:e1005130.

Dearlove BL, Cody AJ, Pascoe B, Méric G, Wilson DJ, Sheppard SK, Daniel J, Sheppard SK. 2016. Rapid host switching in generalist Campylobacter strains erodes the signal for tracing human infections. The ISME journal. 10:721–729.

Didelot X, Bowden R, Wilson DJ, Peto TEA, Crook DW. 2012. Transforming clinical microbiology with bacterial genome sequencing. Nature Reviews Genetics. 13:601–612.

Didelot X, Croucher NJ, Bentley SD, Harris SR, Wilson DJ. 2018. Bayesian inference of ancestral dates on bacterial phylogenetic trees. Nucleic Acids Research. 46:e134.

Didelot X, Fraser C, Gardy J, Colijn C. 2017. Genomic infectious disease epidemiology in partially sampled and ongoing outbreaks. Molecular Biology and Evolution. 34:997–1007.

Didelot X, Gardy J, Colijn C. 2014. Bayesian inference of infectious disease transmission from whole-genome sequence data. Molecular Biology and Evolution. 31:1869–1879.

Didelot X, Kendall M, Xu Y, White PJ, McCarthy N. 2021. Genomic Epidemiology Analysis of Infectious Disease Outbreaks Using TransPhylo. Current Protocols. 1:e60.

Didelot X, Nell S, Yang I, Woltemate S, van der Merwe S, Suerbaum S. 2013. Genomic evolution and transmission of Helicobacter pylori in two South African families. Proceedings of the National Academy of Sciences. 110:13880–13885.

Didelot X, Parkhill J. 2022. A scalable analytical approach from bacterial genomes to epidemiology. Philosophical Transactions of the Royal Society B: Biological Sciences. 377:20210246.

Didelot X, Walker AS, Peto TE, Crook DW, Wilson DJ. 2016. Within-host evolution of bacterial pathogens. Nature Reviews Microbiology. 14:150–162.

Drummond AJ, Nicholls GK, Rodrigo AG, Solomon W. 2002. Estimating mutation parameters, population history and genealogy simultaneously from temporally spaced sequence data. Genetics. 161:1307–1320.

Friel N, Wyse J. 2012. Estimating the evidence–a review. Statistica Neerlandica. 66:288–308.

Gardy JL, Loman NJ. 2018. Towards a genomics-informed, real-time, global pathogen surveillance system. Nature Reviews Genetics. 19:9–20.

Ghafari M, Lumby CK, Weissman DB, Illingworth CJR. 2020. Inferring Transmission Bottleneck Size from Viral Sequence Data Using a Novel Haplotype Reconstruction Method. Journal of Virology. 94:e00014–20.

Green PJ. 1995. Reversible Jump Markov Chain Monte Carlo Computation and Bayesian Model Determination. Biometrika. 82:711–732.

Grenfell BT, Pybus OG, Gog JR, Wood JLN, Daly JM, Mumford JA, Holmes EC. 2004. Unifying the epidemiological and evolutionary dynamics of pathogens. Science. 303:327–332.

Grote A, Earl AM. 2022. Within-host evolution of bacterial pathogens during persistent infection of humans. Current Opinion in Microbiology. 70:102197.

Hall M, Woolhouse M, Rambaut A. 2015. Epidemic Reconstruction in a Phylogenetics Framework: Transmission Trees as Partitions of the Node Set. PLOS Computational Biology. 11:e1004613.

Hall MD, Holden MT, Srisomang P, Mahavanakul W, Wuthiekanun V, Limmathurotsakul D, Fountain K, Parkhill J, Nickerson EK, Peacock SJ, et al. (11 co-authors). 2019. Improved characterisation of MRSA transmission using within-host bacterial sequence diversity. eLife. 8:e46402.

Ho SYW, Shapiro B. 2011. Skyline-plot methods for estimating demographic history from nucleotide sequences. Molecular Ecology Resources. 11:423–434.

Jelley L, Douglas J, Ren X, Winter D, McNeill A, Huang S, French N, Welch D, Hadfield J, de Ligt J, et al. (11 co-authors). 2022. Genomic epidemiology of delta sars-cov-2 during transition from elimination to suppression in aotearoa new zealand. Nature Communications. 13:4035.

Jombart T, Cori A, Didelot X, Cauchemez S, Fraser C, Ferguson N. 2014. Bayesian Reconstruction of Disease Outbreaks by Combining Epidemiologic and Genomic Data. PLoS Computational Biology. 10:e1003457.

Jombart T, Eggo RM, Dodd PJ, Balloux F. 2011. Reconstructing disease outbreaks from genetic data: A graph approach. Heredity. 106:383–390.

Kapli P, Yang Z, Telford MJ. 2020. Phylogenetic tree building in the genomic age. Nature Reviews Genetics. 21:428–444.

Klinkenberg D, Backer JA, Didelot X, Colijn C, Wallinga J. 2017. Simultaneous inference of phylogenetic and transmission trees in infectious disease outbreaks. PLoS Computational Biology. 13:e1005495.

Leitner T. 2019. Phylogenetics in HIV transmission: Taking within-host diversity into account. Current Opinion in HIV and AIDS. 14:181–187.

Lemey P, Rambaut A, Drummond AJ, Suchard MA. 2009. Bayesian phylogeography finds its roots. PLoS computational biology. 5:e1000520.

Lieberman TD, Michel JB, Aingaran M, Potter-Bynoe G, Roux D, Davis MR Jr, Skurnik D, Leiby N, LiPuma JJ, Goldberg JB, et al. (13 co-authors). 2011. Parallel bacterial evolution within multiple patients identifies candidate pathogenicity genes. Nature genetics. 43:1275–1280.

Lloyd-Smith JO, Schreiber SJ, Kopp PE, Getz WM. 2005. Superspreading and the effect of individual variation on disease emergence. Nature. 438:355–359.

Marvig RL, Johansen HK, Molin S, Jelsbak L. 2013. Genome analysis of a transmissible lineage of pseudomonas aeruginosa reveals pathoadaptive mutations and distinct evolutionary paths of hypermutators. PLoS genetics. 9:e1003741.

Mather AE, Reid SWJ, Maskell DJ, Parkhill J, Fookes MC, Harris SR, Brown DJ, Coia JE, Mulvey MR, Gilmour MW. 2013. Distinguishable epidemics of multidrug-resistant Salmonella Typhimurium DT104 in different hosts. Science. 341:1514–1517.

Muller NF, Rasmussen DA, Stadler T. 2017. The Structured Coalescent and Its Approximations. Molecular Biology and Evolution. 34:2970–2981.

Ortiz AT, Kendall M, Storey N, Hatcher J, Dunn H, Roy S, Williams R, Williams C, Goldstein RA, Didelot X, et al. (13 co-authors). 2022. Within-host diversity improves phylogenetic and transmission reconstruction of SARS-CoV-2 outbreaks. bioRxiv. p. 2022.06.07.495142.

Paradis E, Schliep K. 2019. Ape 5.0: An environment for modern phylogenetics and evolutionary analyses in R. Bioinformatics. 35:526–528.

Pybus OG, Charleston MA, Gupta S, Rambaut A, Holmes EC, Harvey PH. 2001. The Epidemic Behavior of the Hepatitis C Virus. Science. 292:2323–2325.

Pybus OG, Rambaut A. 2009. Evolutionary analysis of the dynamics of viral infectious disease. Nature Reviews Genetics. 10:540–550.

Rasmussen DA, Volz EM, Koelle K. 2014. Phylodynamic Inference for Structured Epidemiological Models. PLoS Computational Biology. 10:e1003570.

Rau MH, Marvig RL, Ehrlich GD, Molin S, Jelsbak L. 2012. Deletion and acquisition of genomic content during early stage adaptation of pseudomonas aeruginosa to a human host environment. Environmental microbiology. 14:2200–2211.

Romero-Severson E, Skar H, Bulla I, Albert J, Leitner T. 2014. Timing and order of transmission events is not directly reflected in a pathogen phylogeny. Molecular Biology and Evolution. 31:2472–2482.

Romero-Severson EO, Bulla I, Leitner T. 2016. Phylogenetically resolving epidemiologic linkage. Proceedings of the National Academy of Sciences. 113:2690–2695.

Rossi E, La Rosa R, Bartell JA, Marvig RL, Haagensen JAJ, Sommer LM, Molin S, Johansen HK. 2021. Pseudomonas aeruginosa adaptation and evolution in patients with cystic fibrosis. Nature Reviews Microbiology. 19:331–342.

Sagulenko P, Puller V, Neher RA. 2018. TreeTime: Maximum-likelihood phylodynamic analysis. Virus Evolution. 4:vex042.

Spencer SEF. 2021. Accelerating adaptation in the adaptive Metropolis-Hastings random walk algorithm. Australian & New Zealand Journal of Statistics. 63:468–484.

Suchard MA, Lemey P, Baele G, Ayres DL, Drummond AJ, Rambaut A. 2018. Bayesian phylogenetic and phylodynamic data integration using BEAST 1.10. Virus Evolution. 4:vey016.

Tartakovsky AG, Moustakides GV. 2010. State-of-the-art in Bayesian changepoint detection. Sequential Analysis. 29:125–145.

Tonkin-Hill G, Ling C, Chaguza C, Salter SJ, Hinfonthong P, Nikolaou E, Tate N, Pastusiak A, Turner C, Chewapreecha C, et al. (15 co-authors). 2022. Pneumococcal within-host diversity during colonization, transmission and treatment. Nature Microbiology. 7:1791–1804.

van Dorp L, Wang Q, Shaw LP, Acman M, Brynildsrud OB, Eldholm V, Wang R, Gao H, Yin Y, Chen H, et al. (15 co-authors). 2019. Rapid phenotypic evolution in multidrug-resistant Klebsiella pneumoniae hospital outbreak strains. Microbial Genomics. 5:e000263.

Volz EM, Frost SDW. 2017. Scalable relaxed clock phylogenetic dating. Virus Evolution. 3:vex025.

Volz EM, Koelle K, Bedford T. 2013. Viral Phylodynamics. PLoS Computational Biology. 9:e1002947.

Wallinga J, Teunis P. 2004. Different Epidemic Curves for Severe Acute Respiratory Syndrome Reveal Similar Impacts of Control Measures. American Journal of Epidemiology. 160:509–516.

Wymant C, Hall M, Ratmann O, Bonsall D, Golubchik T, de Cesare M, Gall A, Cornelissen M, Fraser C, STOP-HCV Consortium, et al. (12 co-authors). 2018. PHYLOSCANNER: Inferring Transmission from Within- and Between-Host Pathogen Genetic Diversity. Molecular Biology and Evolution. 35:719–733.

Xu Y, Stockdale JE, Naidu V, Hatherell H, Stimson J, Stagg HR, Abubakar I, Colijn C. 2020. Transmission analysis of a large tuberculosis outbreak in London: A mathematical modelling study using genomic data. Microbial Genomics. 6:e000450.

Yang L, Jelsbak L, Marvig RL, Damkiær S, Workman CT, Rau MH, Hansen SK, Folkesson A, Johansen HK, Ciofu O, et al. (13 co-authors). 2011. Evolutionary dynamics of bacteria in a human host environment. Proceedings of the National Academy of Sciences. 108:7481–7486.

Yang S, Hemarajata P, Hindler J, Li F, Adisetiyo H, Aldrovandi G, Sebra R, Kasarskis A, MacCannell D, Didelot X, et al. (13 co-authors). 2017. Evolution and Transmission of Carbapenem-Resistant Klebsiella pneumoniae Expressing the bla_OXA-232_ Gene During an Institutional Outbreak Associated With Endoscopic Retrograde Cholangiopancreatography. Clinical Infectious Diseases. 64:894–901.

Yang Z, Rannala B. 2012. Molecular phylogenetics: Principles and practice. Nature Reviews Genetics. 13:303–314.

Young BC, Golubchik T, Batty EM, Fung R, Larner-Svensson H, Votintseva AA, Miller RR, Godwin H, Knox K, Everitt RG, et al. (21 co-authors). 2012. Evolutionary dynamics of Staphylococcus aureus during progression from carriage to disease. Proceedings of the National Academy of Sciences. 109:4550–4555.

