## Supplementary Material for "Inference of infectious disease transmission using multiple genomes per host"

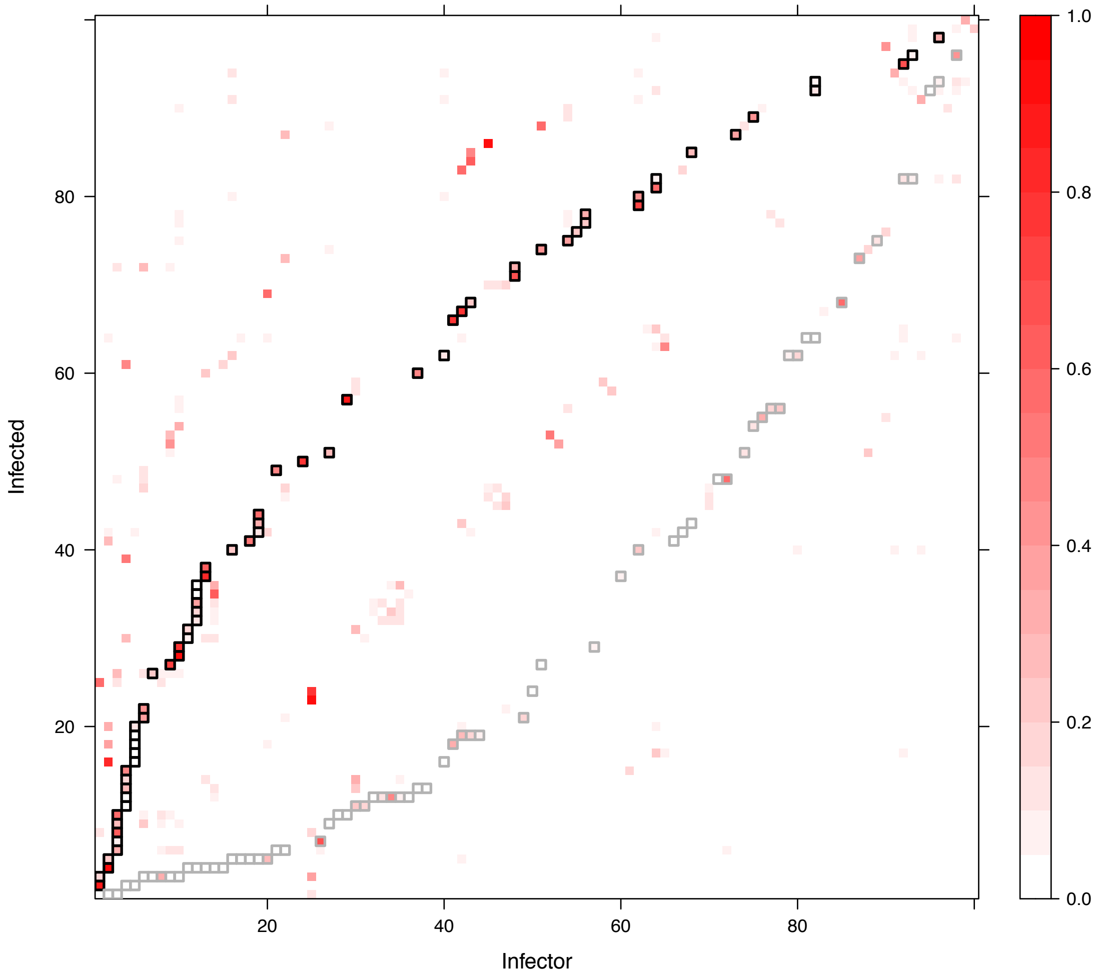


Figure S1: Posterior probability estimates of transmissions from an infector (row) to an infected host (column) for a simulated dataset with one observation per host. The black squares show the true transmissions in the simulated dataset. The gray squares show the reverse relationship, switching the true infector and infected hosts.

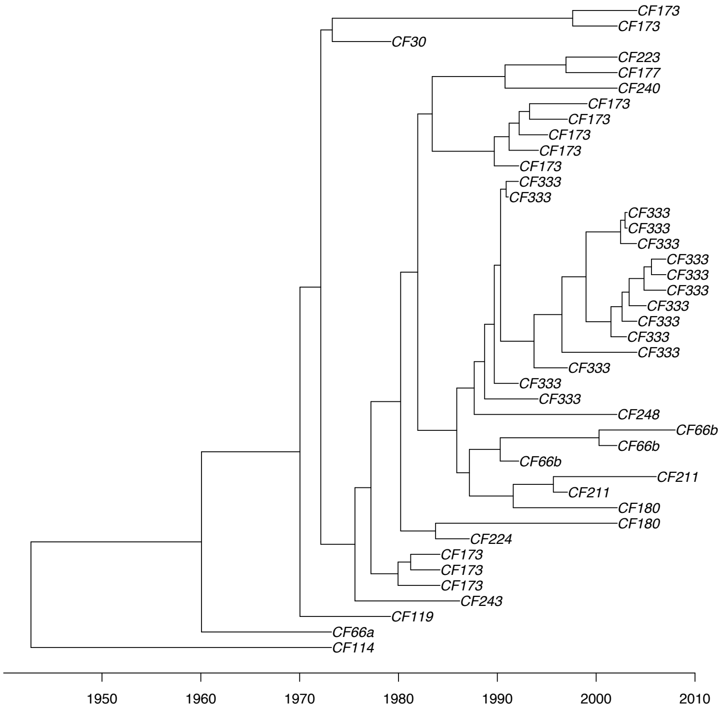


Figure S2: Dated tree used as input of the *P. aeruginosa* analysis.

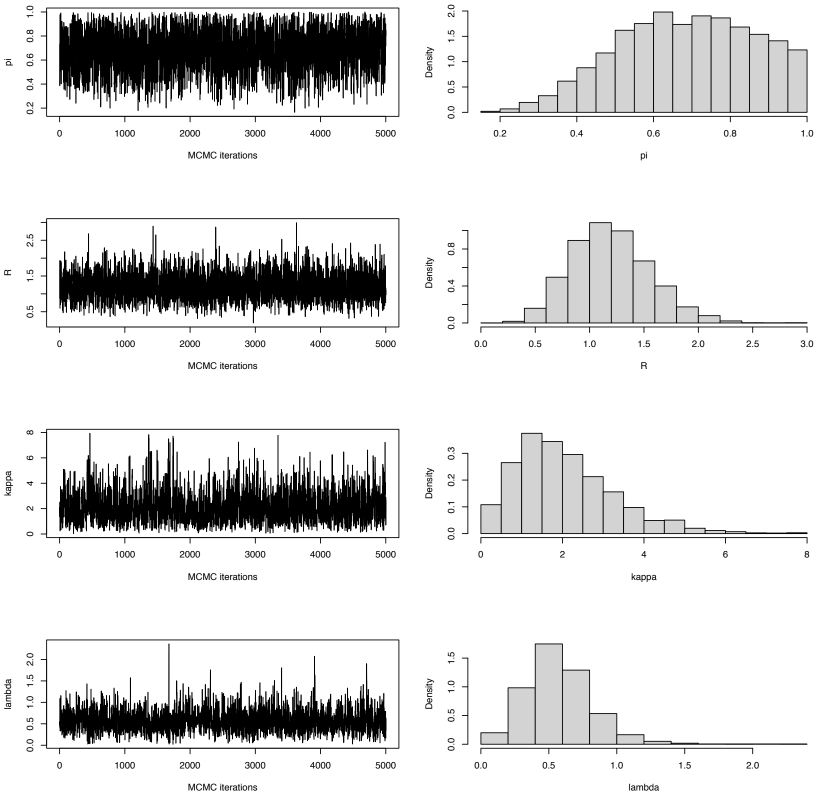


Figure S3: Parameter estimates in the *P. aeruginosa* analysis.

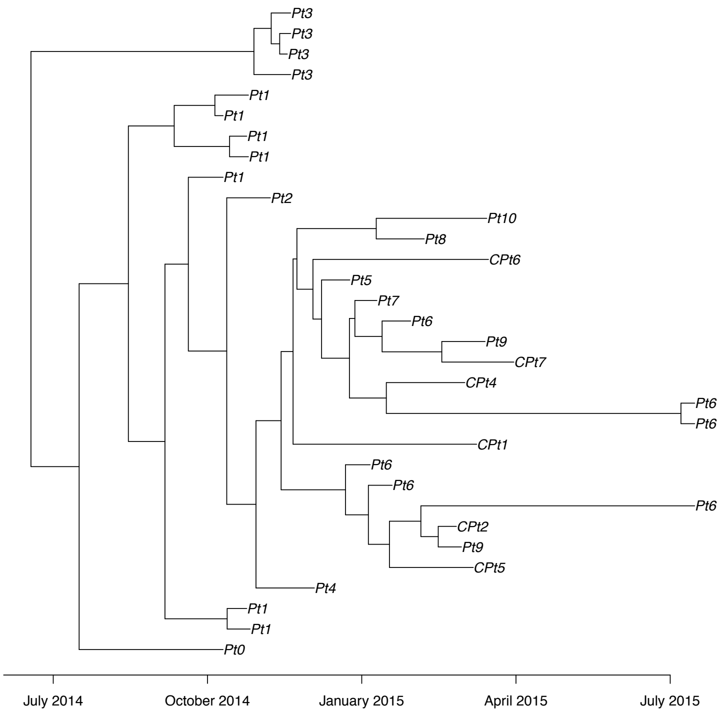


Figure S4: Dated tree used as input of the *K. pneumoniae* analysis.

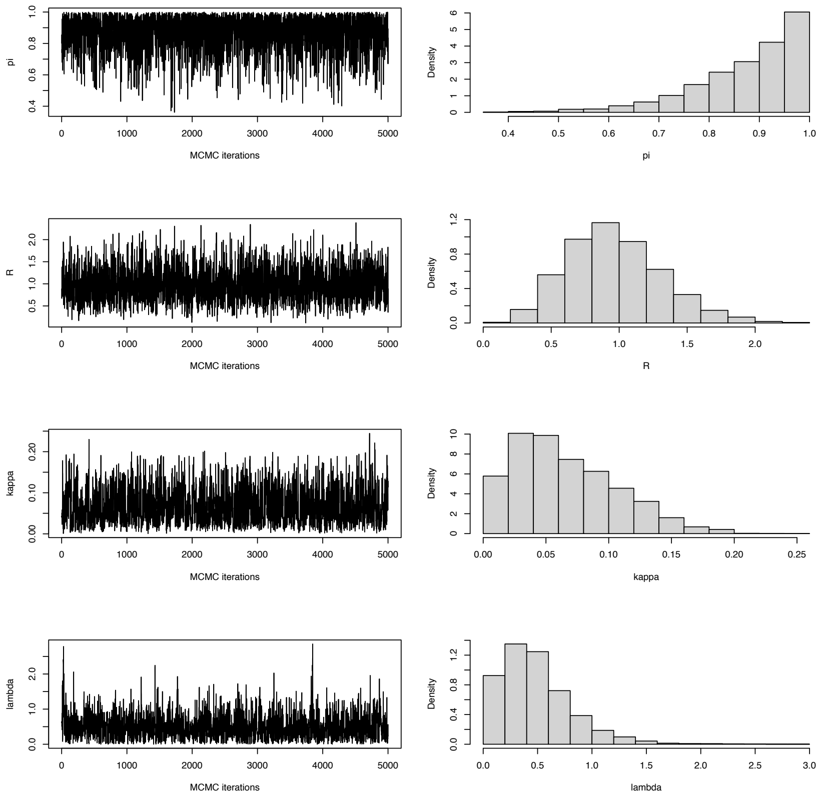


Figure S5: Parameter estimates in the *K. pneumoniae* analysis.

**Algorithm S1** Simulation of a transmission tree in an ongoing outbreak scenario with cut-off time $T$

Initialise root time $x^1$, $j \leftarrow 1$, $h \leftarrow 1$.
**while** $j \leq h$ **do**
With probability $\frac{\varpi(x^j)}{1-\omega(x^j)}$ set $S^j \leftarrow 1$, otherwise set $S^j \leftarrow 0$.
**if** $S^j = 1$ **then**
Sample primary observation time from $\sigma(t - x^j)$.
Sample the number of secondary observations according to $\beta(b)$ and their times according to $\rho(\tau_{1:b})$.
Update vectors $y$, $z$, $H_y$, and $H_z$, censoring any observation times exceeding $T$.
Sample number of included offspring $d^j \sim p(d \mid x^j)$.
**else**
Sample number of included offspring $d^j \sim p(d \mid x^j)$, $d > 0$.
**end if**
**for** $g = 1, ..., d^j$ **do**
Set $h \leftarrow h + 1$ and set $A^h = j$.
Sample infection time

$$x^h \sim \frac{(1 - \omega(t))\gamma(t - x^j)}{1 - \bar{\omega}(x^j)}, \qquad x^j < t < T.$$

**end for**
Set $j \leftarrow j + 1$.
**end while**

**Algorithm S2** Simulation of a transmission tree in an ongoing outbreak scenario with maximum number of observed hosts $N$

Initialise root time $x^1$, $j \leftarrow 1$, $h \leftarrow 1$, $n \leftarrow 0$, $T' \leftarrow \infty$.
**while** $j \leq h$ **do**
  with probability $\pi$ set $S^j \leftarrow 1$, otherwise set $S^j \leftarrow 0$.
  **if** $S^j = 1$ **then**
    Propose primary observation time from $\sigma(t - x^j)$.
    Update number of observed hosts $n \leftarrow n + 1$.
    **if** $n > N + 1$ **then**
      Set $T'$ as the time of the $(N+1)$'th primary observation, ordered earliest to latest.
    **end if**
    Propose number of secondary observations according to $\beta(b)$ and their times according to $\rho(\tau_{1:b})$.
    Update vectors $y$, $z$, $H_y$, and $H_z$, censoring any observations from hosts with a primary observation time later than $T'$.
  **end if**
  Sample number of offspring $k^j \sim \alpha(k)$.
  **for** $g = 1, ..., k^j$ **do**
    Propose infection time $t' \sim \gamma(t - x^j)$.
    **if** $t' < T'$ **then**
      Set $h \leftarrow h + 1$ and set $A^h = j$.
      Set $x^h \leftarrow t'$.
    **end if**
  **end for**
**end while**
Set $T$ as a time between the $N$'th and $(N+1)$'th primary observation.
Censor any observations from hosts with a primary observation time later than $T$, updating $y$, $z$, $H_y$, and $H_z$ accordingly.
Discard excluded hosts, updating $x$ and $A$ accordingly.

**Algorithm S3** Simulation of a transmission tree in an ongoing outbreak scenario with maximum number of observations $N$

Initialise root time $x^1$, $j \leftarrow 1$, $h \leftarrow 1$, $n \leftarrow 0$, $T' \leftarrow \infty$.
**while** $j \leq h$ **do**
  with probability $\pi$ set $S^j \leftarrow 1$, otherwise set $S^j \leftarrow 0$.
  **if** $S^j = 1$ **then**
    Propose primary observation time from $\sigma(t - x^j)$.
    Propose number of secondary observations according to $\beta(b)$ and their times according to $\rho(\tau_{1:b})$.
    Update vectors $y$, $z$, $H_y$, and $H_z$, censoring any observation times exceeding $T'$.
    Update number of observations $n$.
    **if** $n > N + 1$ **then**
      Set $T'$ as the time of the $(N + 1)$'th observation, ordered earliest to latest.
    **end if**
  **end if**
  Sample number of offspring $k^j \sim \alpha(k)$.
  **for** $g = 1, ..., k^j$ **do**
    Propose infection time $t' \sim \gamma(t - x^j)$.
    **if** $t' < T'$ **then**
      Set $h \leftarrow h + 1$ and set $A^h = j$.
      Set $x^h \leftarrow t'$.
    **end if**
  **end for**
**end while**
Set $T$ as a time between the $N$'th and $(N + 1)$'th observation.
Censor any observations later than $T$, updating $y$, $z$, $H_y$, and $H_z$ accordingly.
Discard excluded hosts, updating $x$ and $A$ accordingly.

**Algorithm S4** Simulation of a host subtree

Initialise $t \leftarrow v_j^M$; $L \leftarrow 1$; $m \leftarrow M$; $r \leftarrow 0$

**while** $t > x^j$ **do**

  Propose coalescence time

$$u_j^* = x^j + \frac{1}{\lambda}\left((1-\nu)^{\frac{\lambda}{\binom{L}{2}}}(\kappa + \lambda(t - x^j)) - \kappa\right)$$

  **if** $u_j^* > v_j^{m-1}$ **then**

    Set coalescence time $r \leftarrow r+1$; $u_j^r \leftarrow u_j^*$; $L \leftarrow L-1$; $t \leftarrow u_j^*$

  **else**

    $m \leftarrow m-1$; $L \leftarrow L+1$; $t \leftarrow v_j^m$

  **end if**

**end while**

**for** $n = 1, ..., r$ **do**

  Reverse coalescence times $u_j^n \leftarrow u_j^{r+1-n}$

**end for**
